# RUNX1/RUNX1T1 controls alternative splicing in the t(8;21)-positive acute myeloid leukemia cells

**DOI:** 10.1101/628040

**Authors:** Vasily Grinev, Ilia Ilyushonak, Richard Clough, Sirintra Nakjang, Job Smink, Natalia Martinez-Soria, Tatsiana Ramanouskaya, Constanze Bonifer, Olaf Heidenreich

**Author notes:** Correspondence (V.G.), (O.H.).

## Abstract

The fusion oncogene *RUNX1/RUNX1T1* encodes an aberrant transcription factor, which plays a key role in the initiation and maintenance of the t(8;21)-positive acute myeloid leukemia. Here we show that this oncogene is a regulator of the alternative RNA splicing for a sub-set of genes in the leukemia cells. We found two primary mechanisms underlying changes in the production of RNA isoforms: (i) RUNX1/RUNX1T1-mediated regulation of alternative transcription start sites selection in target genes, and (ii) direct or indirect control of the expression of the genes encoding splicing factors. The first mechanism leads to the expression of RNA isoforms with alternative structure of the 5’-UTR regions. The second mechanism generates alternative transcripts with new junctions between internal cassettes and constitutive exons. We also show that RUNX1/RUNX1T1-mediated differential splicing affects several functional groups of genes and produces proteins with unique conserved domain structures. In summary, this study reveals a novel layer of RUNX1/RUNX1T1-dependent transcriptome organization in t(8;21)-positive acute myeloid leukemia.

## INTRODUCTION

The fusion oncogene *RUNX1/RUNX1T1* is a result of the nonhomologous reciprocal translocation t(8;21), which involves the *RUNX1* gene on chromosome 21 and the *RUNX1T1* gene on chromosome 8 (Miyoshi et al., 1991). When expressed in haematopoietic cells, the fusion protein occupies more than 4000 genomic sites (Gardini et al., 2008; Maiques-Diaz et al., 2012; Mandoli et al., 2016; Li et al., 2016; Trombly et al., 2015) and forms transcription regulatory complexes by recruiting co-factors (Corpora et al., 2010; Liu et al., 2006; Park et al., 2009; Sun et al., 2013). These complexes trigger a local remodeling of chromatin of a wide range of genes (Ptasinska et al., 2012; Wang et al., 2011) and, thereby, affect their expression (Gardini et al., 2008; Li et al., 2016). In turn, the change of target gene expression leads to a block of cell differentiation (Ptasinska et al., 2014), enhancement of self-renewal (Martinez-Soria et al., 2018; Ptasinska et al., 2014), modulation of the apoptosis (Mandoli et al., 2016; Spirin et al., 2014) and, eventually, to malignant transformation of the t(8;21)-positive cells. However, despite this increasing comprehension of the molecular function of the *RUNX1/RUNX1T1*, its impact on the leukaemic transcriptome is still only incompletely understood.

Alternative splicing increases the transcriptomic and proteomic complexity by generating distinct isoforms with different roles and functions from a single gene (Baralle and Giudice, 2017). When dysregulated, alternative splicing can contribute to tumorigenesis (Kozlovski et al., 2017; Singh and Eyras, 2017). Missplicing of genes has been found in multiple malignancies including carcinomas, neuroblastoma, chronic lymphoid leukemia and acute myeloid leukaemia (AML) (Adamia et al., 2014). Mutations in spliceosome factor genes such as *SF3B1, SRSF2* or *U2AF* are frequently found in myelodysplastic syndrome (Qiu et al., 2016) and are currently intensively examined for their therapeutic relevance (Lee and Abdel-Wahab, 2016; Lee et al., 2016). In addition, alternative splicing is also regulated by epigenetic marks (Hnilicová et al., 2011; Zhou et al., 2014). Histone modifications and DNA methylation can control exon usage by controlling the elongation speed of RNA polymerase II and, consequently, the choice of splice sites. Furthermore, they may affect the recruitment of splicing factors to the chromatin through adaptors such as CHD1. Moreover, epigenetic marks have been found to regulate the activity of alternative transcriptional start sites (TSSs) including cryptic ones in the genome (Brocks et al., 2017). A potential role of leukaemic fusion genes encoding epigenetic modulators such as RUNX1/RUNX1T1 has not yet been established for the regulation of alternative splicing.

To address this question, we performed perturbation experiments in t(8;21)-positive AML cells and examined the impact of RUNX1/RUNX1T1 knockdown on the gene expression and RNA splicing at a global level. Our results demonstrate that *RUNX1/RUNX1T1* downregulation affects splicing of both direct and indirect target genes. Mechanistically, changes in the production of RNA isoforms are implemented via (i) direct RUNX1/RUNX1T1-mediated control of alternative transcription start sites selection in target genes, and (ii) direct or indirect control of the expression of the genes encoding splicing factors. Our *in silico* modeling and experimental results indicate that RUNX1/RUNX1T1-mediated differential splicing affects conserved domain structures in proteins ultimately modulating leukaemia-relevant processes such as nucleotide metabolism, cell adhesion and cell differentiation.

In conclusion, our results show that the fusion oncogene *RUNX1/RUNX1T1* controls alternative splicing in t(8;21)-positive AML cells. This reveals a new layer of complexity in the organization of t(8;21)-positive AML cells and sets a paradigm for the role of fusion gene-encoded transcription factors in RNA processing.

## RESULTS

### Change in the expression of the fusion oncogene *RUNX1/RUNX1T1* leads to differential splicing in the transcriptome of Kasumi-1 cells

We initially analyzed published data sets (Table S1) for a potential association between RUNX1/RUNX1T1 and RNA processing. Functional annotation analysis uncovered a highly significant enrichment of RUNX1/RUNX1T1 binding at gene loci known to be subject to alternative splicing and splice variants (Table S2, FDR-adjusted p-value, or q-value, 3.98 × 10^−23^ and 1.48 × 10^−12^, respectively). These findings are in line with the observation that RUNX1/RUNX1T1 may interact with several splicing factors implying a direct link between RUNX1/RUNX1T1-regulated transcription and RNA processing (Mandoli et al., 2016). Furthermore, RUNX1/RUNX1T1 binding sites are often present within the genes encoding splicing-associated factors or in their immediate vicinity. The list of such genes includes both classical splicing regulatory genes (HNRNPA1, RBM28, SF3A3, SRSF5, etc.) and genes encoding the components of nuclear speckles (DDX42, GLI3, SARNP, etc.), which are considered the area of intensive splicing (Cardoso et al., 2012; Lamond et al., 2003). Moreover, gene set enrichment analysis suggests that knockdown of *RUNX1/RUNX1T1* is associated with impaired mRNA processing and, in particular, splicing pathways (Figures 1A, S1A). These findings strongly support a regulatory role of this leukaemic fusion protein in mRNA splicing and predict changes in splicing pattern in response to perturbing RUNX1/RUNX1T1 activity.

To identify such differential splicing events, we performed RNA-Seq analysis in the t(8;21)-positive AML cell line Kasumi-1 following *RUNX1/RUNX1T1* knockdown. Knockdown was achieved by electroporation with the siRNA siRR targeting the fusion site of the *RUNX1/RUNX1T1* mRNA, while a mismatch short interfering RNA, siMM, served as a control (Figure 1B) (Heidenreich et al., 2003). The differential splicing events were detected at the level of both exon usage and exon-exon junctions (EEJs) using DEXSeq, limma/diffSplice and JunctionSeq (Anders et al., 2012; Hartley et al., 2016; Ritchie et al., 2015). The combined use of these complementing algorithms detected a comprehensive set of differential splicing events in dependence on the RUNX1/RUNX1T1 status.

**Figure 1.**
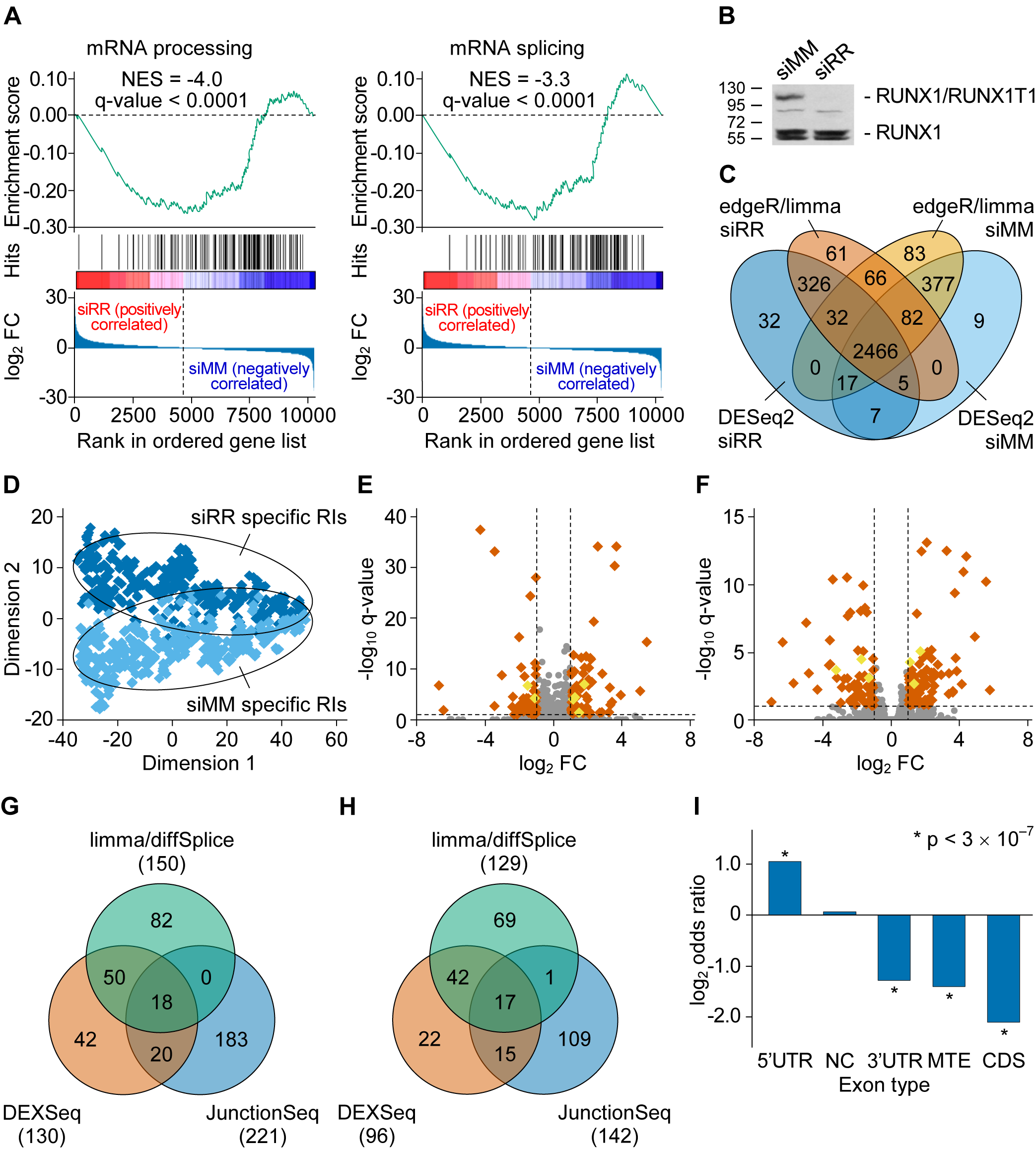
Knockdown of the fusion oncogene *RUNX1/RUNX1T1* leads to differential usage of a subset of exons in the transcriptome of Kasumi-1 cells. (A) The GSEA enrichment plots for the Reactome mRNA processing and splicing pathways. The normalized enrichment score (NES) and its statistical significance (q-value) are shown in the each enrichment plot. (B) Western blot of RUNX1 and RUNX1/RUNX1T1 protein levels in Kasumi-1 cells following RUNX1/RUNX1T1 knockdown. (C) Distribution of RIs between the transcriptomes of the siRR- and siMM-treated leukemia cells. The RIs were detected using the DESeq2 and edgeR/limma algorithms. (D) Multidimensional data reduction using t-distributed stochastic neighbor embedding allows to clearly separating the siRR- and siMM-specific RIs. This separation is based on the expression data of RIs. (E) The DEXSeq algorithm identified 130 diffUEs distributed over 96 individual genes. These exons include 6 RIs indicated with yellow rhombus. (F) The limma/diffSplice algorithm identified 150 diffUEs distributed over 129 individual genes. These exons include 6 RIs indicated with yellow rhombus. (G) The Venn diagram demonstrates the degree of overlap between the lists of diffUEs identified by the DEXSeq, limma/diffSplice and JunctionSeq algorithms. (H) The Venn diagram demonstrates the degree of overlap between the lists of genes with diffUEs identified by the DEXSeq, limma/diffSplice and JunctionSeq algorithms. (I) Classification of diffUEs according to the functional type of exons. Genomic coordinates of the reference exons were extracted from Ensembl models of the human genes. diffUEs were intersected with reference exons and the overlaps counted. Fisher’s exact test was performed to detect over- or under-represented exon types among the diffUEs against non-diffUEs.

Using our pipeline, we analysed 511,363 non-overlapping bins of human canonical exons (CEs) annotated in Ensembl (Zerbino et al., 2018). This exonic list was extended by 3540 non-overlapping bins of retained introns (RIs) detected in 2401 genes of the Kasumi-1 cells (Table S3). Of all retained introns, 419 and 469 were exclusively expressed in either RUNX1/RUNX1T1 knockdown or control leukemia cells (Figures 1C, D, S1B). Both summarized read and assembled transcriptome data indicated for genes containing such condition-specific retained introns a trend towards changed expression after *RUNX1/RUNX1T1* knockdown (Figures S1D-S1H).

With the above-mentioned combined list of bins, we identified 395 differentially used exons (diffUEs) in the Kasumi-1 transcriptome following *RUNX1/RUNX1T1* knockdown (Figures 1E, 1F, S1I, Table S4). The list of the identified diffUEs included 387 canonical exons and 8 retained introns distributed over 275 individual genes (Figures 1G, 1H). These diffUEs were significantly enriched for 5’-UTR exons and were depleted of CDS, 3’-UTR and multi-type exons (MTE; can be non-coding, 5’-UTR, 5’-UTR/CDS, CDS, CDS/3’-UTR and/or 3’-UTR exon). At the same time, no preference was found among the exons of non-coding RNAs (Figure 1I).

At the level of EEJs, we detected 177 differential EEJs (diffEEJs) upon *RUNX1/RUNX1T1* knockdown that were distributed over 150 genes (Figures 2A-C, S1A, Table S5). These diffEEJs belong to the main modes of alternative splicing (Figures 2D, E) and associated with all functional types of exons (Figure 2F). Notably, use of two different algorithms diffSplice and JunctionSeq for the analysis of EEJs produced only minimally overlapping results (Figures 2B, C). This is due to the differences in the pre-processing, normalization and transformation of the data, as well as in the use of different statistical models at the step of identification of diffEEJs by the algorithms. Nevertheless, diffEEJs identified by both diffSplice and JunctionSeq were confirmed using qPCR (Table S6). In contrast to genes with condition-specific retained introns, only 55 of 378 genes with diffUEs or diffEEJs showed changes in total transcript levels upon RUNX1/RUNX1T1 knockdown (Figures S2B, C).

**Figure 2.**
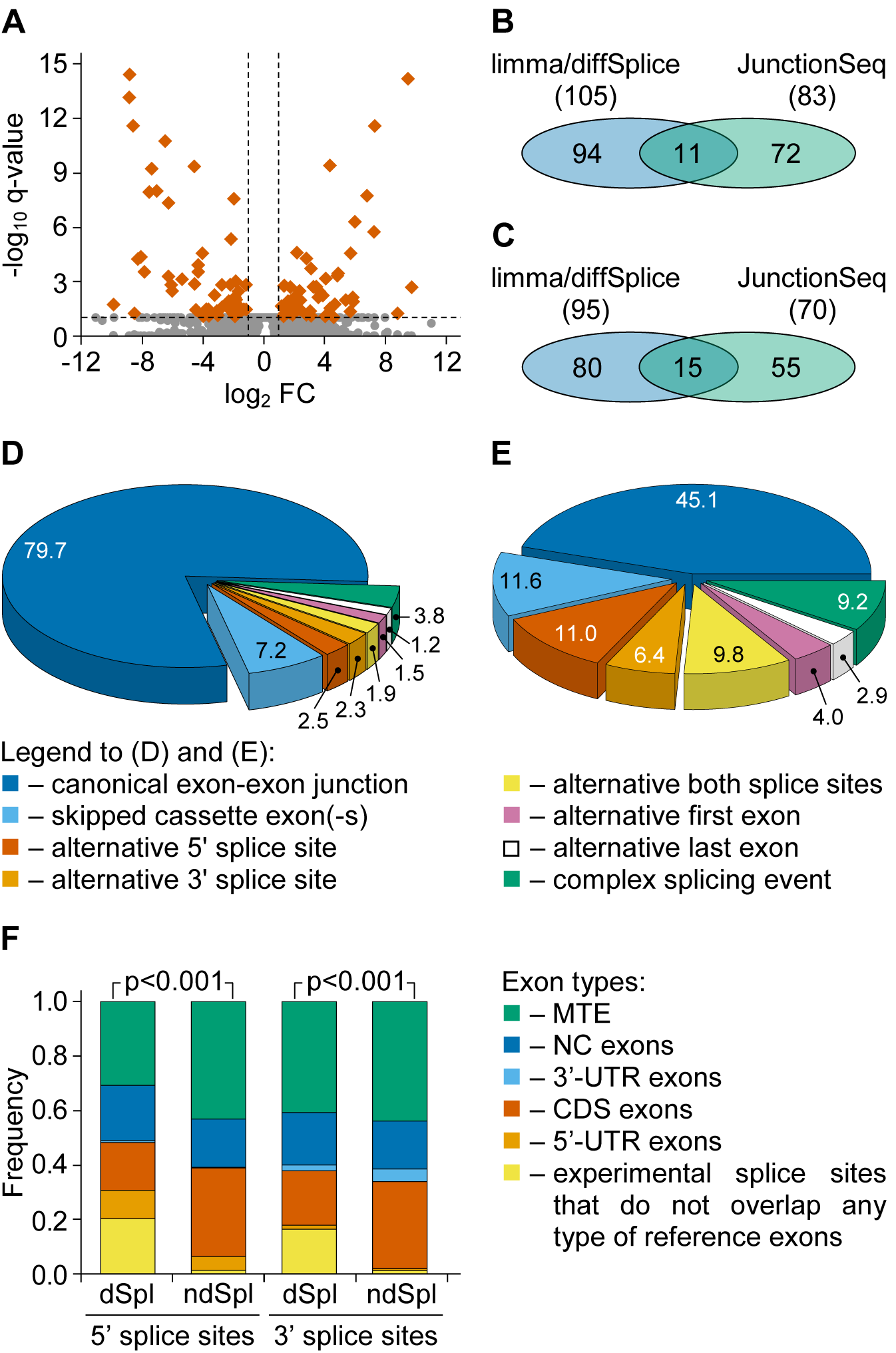
Knockdown of the fusion oncogene *RUNX1/RUNX1T1* leads to differential usage of a subset of EEJs in the transcriptome of Kasumi-1 cells. (A) The limma/diffSplice algorithm identified 105 diffEEJs distributed over 95 individual genes. (B) The Venn diagram demonstrates the depth of overlap between the lists of diffEEJs identified by the limma/diffSplice and JunctionSeq algorithms. (C) The Venn diagram demonstrates the degree of overlap between the lists of genes with diffEEJs identified by the limma/diffSplice and JunctionSeq algorithms. (D) and (E) Classification of non-diffEEJs (D) and diffEEJs (E) according to the modes of alternative splicing. Classification procedure was performed using Ensembl-based models of hypothetical “non-alternative” precursor of RNAs of human genes. Numbers show the percentage of EEJs assigned to a particular mode of splicing. (F) There is a statistically significant (according to the χ^2^ test) shift in the distribution of the splice sites of the diffEEJs (dSpl) and non-diffEEJs (ndSpl) among the various functional types of exons. To produce this picture, genomic coordinates and functional annotations of exons were retrieved from Ensembl models of the human genes. Next, experimentally identified splice sites were intersected with splice sites of reference exons and finally overlaps were counted.

Together, these results clearly indicate that perturbation of RUNX1/RUNX1T1 expression leads to differential splicing of the leukaemic transcriptome.

### Epigenetic marks affect alternative splicing

Binding of RUNX1/RUNX1T1 at specific genomic sites affects the local chromatin status, including changes in histone modifications and chromatin accessibility (Maiques-Diaz et al., 2012; Ptasinska et al., 2012; Wang et al., 2011) (Figures 3A, upper panel, H3K9Ac data). In turn, local remodeling of chromatin leads to changes in the transcription rate, which can affect alternative splicing (Nieto Moreno et al., 2015; Ramanouskaya et al., 2017). Furthermore, we observed redistributions of the RNA polymerase II peaks in siRR-comparing to siMM-treated leukemia cells (Figures 3A, upper panel, RNA polymerase II data).

Splice sites of diffEEJs are significantly enriched in the regions of the open chromatin (marked by DNase I hypersensitivity (HS) sites and sites of H3K9Ac histone modification) and slowed RNA polymerase II elongation compared to the splice sites of non-diffEEJs (Figure 3B). We also observed that the alternative 5’ splice sites (but not 3’ splice sites) of diffEEJs are more proximal to open chromatin regions and sites bound by H3K9Ac and Pol II then canonical ones (Figure 3C). Similarly, alternative 5’ splice sites showed also a trend towards closer proximity to RUNX1/RUNX1T1 sites. These data suggest that regions of H3K9Ac histone modification and RNA polymerase II occupation, which cover the splice sites of diffEEJs, are under the control of RUNX1/RUNX1T1 (Figure 3A, bottom panel).

**Figure 3.**
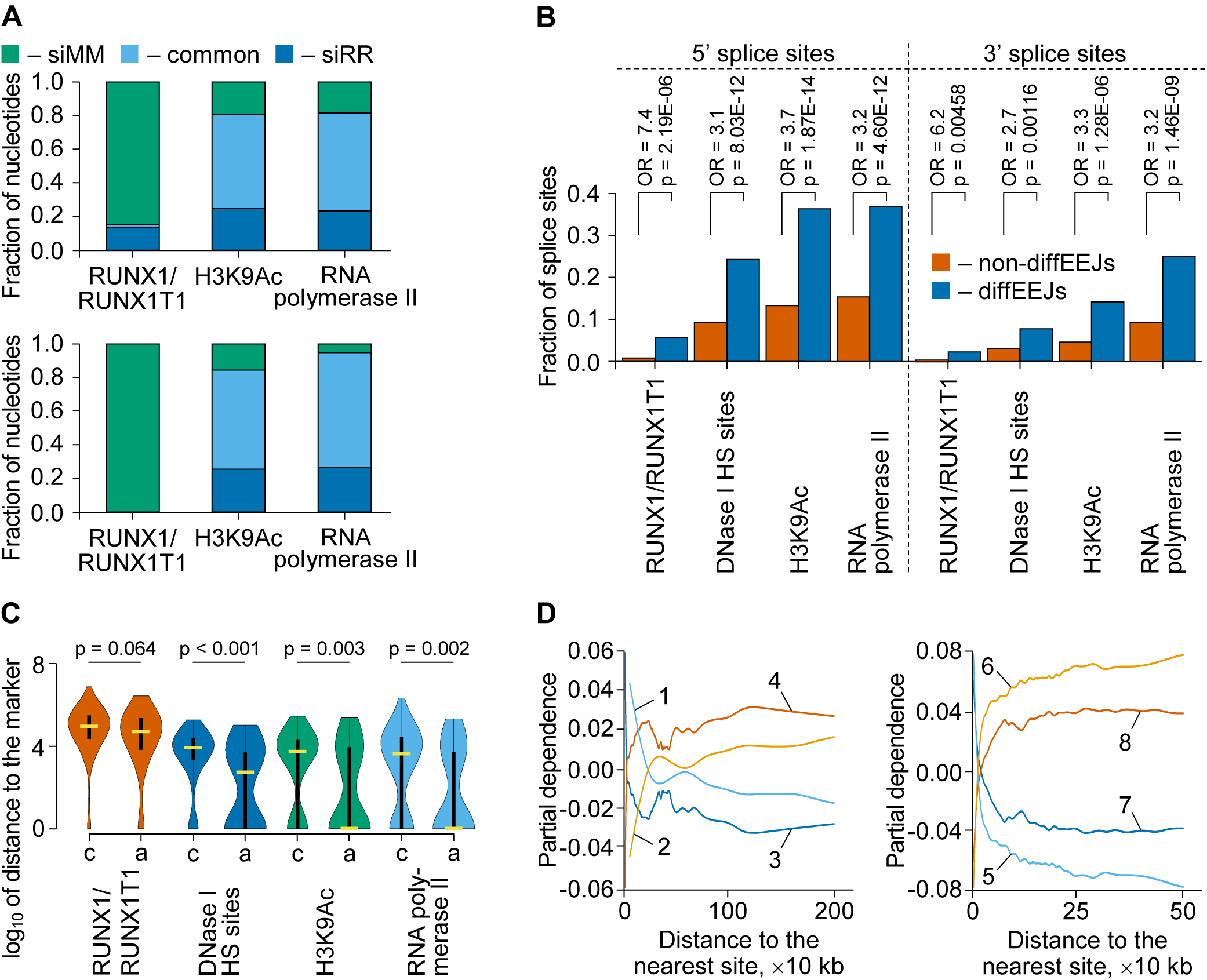
*RUNX1/RUNX1T1*-dependent differential splicing demonstrates a close association with open chromatin. (A) The knockdown of the fusion oncogene *RUNX1/RUNX1T1* affects epigenetic landscape in Kasumi-1 cells. On the upper panel, all RUNX1/RUNX1T1, H3K9Ac or RNA polymerase II bound genomic intervals from siMM- and siRR-treated cells were disjointed into unique non-overlapping bins. These bins were clustered into three groups: siMM-specific (siMM; these bins were occupied by indicated protein only in siMM-treated cells), siRR-specific (siRR; these bins were identified only in siRR-treated cells) and non-specific (common; these bins were bound by indicated protein in both type of cells). Finally, fraction of nucleotides in each group of bins and for each protein was calculated and plotted. On the bottom panel, only ChIP-Seq peaks overlapping of the 5’ splice sites of diffEEJs were selected for subsequent processing and plotting as described in the upper panel. Very similar distributions were observed with 3’ splice sites of the diffEEJs. (B) A significant fraction of the splice sites of diffEEJs falls into open regions of the chromatin of Kasumi-1 cells. According to Fisher’s exact test, the splice sites of diffEEJs demonstrate a much stronger association (in terms of odds ratio, OR) with marks of open chromatin compared to the splice sites of non-diffEEJs. This picture is based on data obtained from siMM-treated cells but very similar results were observed with data from siRR-treated ones. (C) Positional relationships between the canonical (c) and alternative (a) 5’ splice sites of diffEEJs and marks of open chromatin. Grouping of 5’ splice sites into two classes was performed using Ensembl-based models of hypothetical “non-alternative” precursor of RNAs of human genes. According to Mann-Whitney U test, the alternative 5’ splice sites are located significantly closer to the DNase I HS sites, H3K9Ac histone modifications and RNA polymerase II peaks then canonical ones. On each violin plot, the yellow horizontal thick line represents the median of distances distribution, the black vertical rectangular box shows the interquartile range, and the black vertical line is the 95% confidential interval. This picture is based on data obtained from siMM-treated cells but very similar results were observed with data from siRR-treated ones. It is noteworthy that no significant differences were found between the canonical and alternative 3’ splice sites of diffEEJs. (D) Effect of the proximity to RUNX1/RUNX1T1 binding sites (lines 1 and 2), RNA polymerase II peaks (lines 3 and 4), H3K9Ac histone modifications (lines 5 and 6) and DNase I HS sites (lines 7 and 8) on the probability of the diffEEJs (lines 1, 3, 5 and 7) and non-diffEEJs (lines 2, 4, 6 and 8) in the transcriptome of Kasumi-1 cells. These partial dependence plots are based on the results from 1000 independent runs of the random forest meta-classifier and 1000 classification trees per random forest per run.

All of the above findings indicate a close association between chromatin state and differential splicing following *RUNX1/RUNX1T1* knockdown. However, the epigenetic landscape affects alternative splicing only in cooperation with sequence features of pre-mRNA (Luco et al., 2011). To integrate epigenetic data with multivariate sequence data and to calculate partial dependence for each epigenetic mark, we used a predictive model based on the random forest of decision trees (see Supplemental Experimental Procedures for further details). Partial dependence shows the adjusted effect of a given feature on the probability whether a splicing event belongs to the differential or non-differential class in the context of the whole set of other features (Friedman, 2001). This analysis demonstrates that differential splicing is promoted by proximity to RUNX1/RUNX1T1, RNA polymerase II and H3K9Ac sites as well as DNase I HS sites (Figures 3D) and highlights the influence of the chromatin state on differential splicing following RUNX1/RUNX1T1 knockdown.

### Delayed splicing is sensitive to regulation by RUNX1/RUNX1T1

The majority of splicing events are co-transcriptional. However, in some cases of regulated alternative splicing, intron removal can be delayed until the release of pre-mRNA from the site of transcription (Boutz et al., 2015; Braun et al., 2017; Vargas et al., 2011). With this in mind, we analysed the nascent RNA in Kasumi-1 cells for EEJs that were shared between nascent and total RNA (Figure 4A, B). These EEJs were grouped into two classes. The first class (non-diffEEJs) included the nascent EEJs which did not become differential EEJs at the level of total RNA. In contrast, the second class (diffEEJs) collected events that were identified as diffEEJs at the level of total RNA.

**Figure 4.**
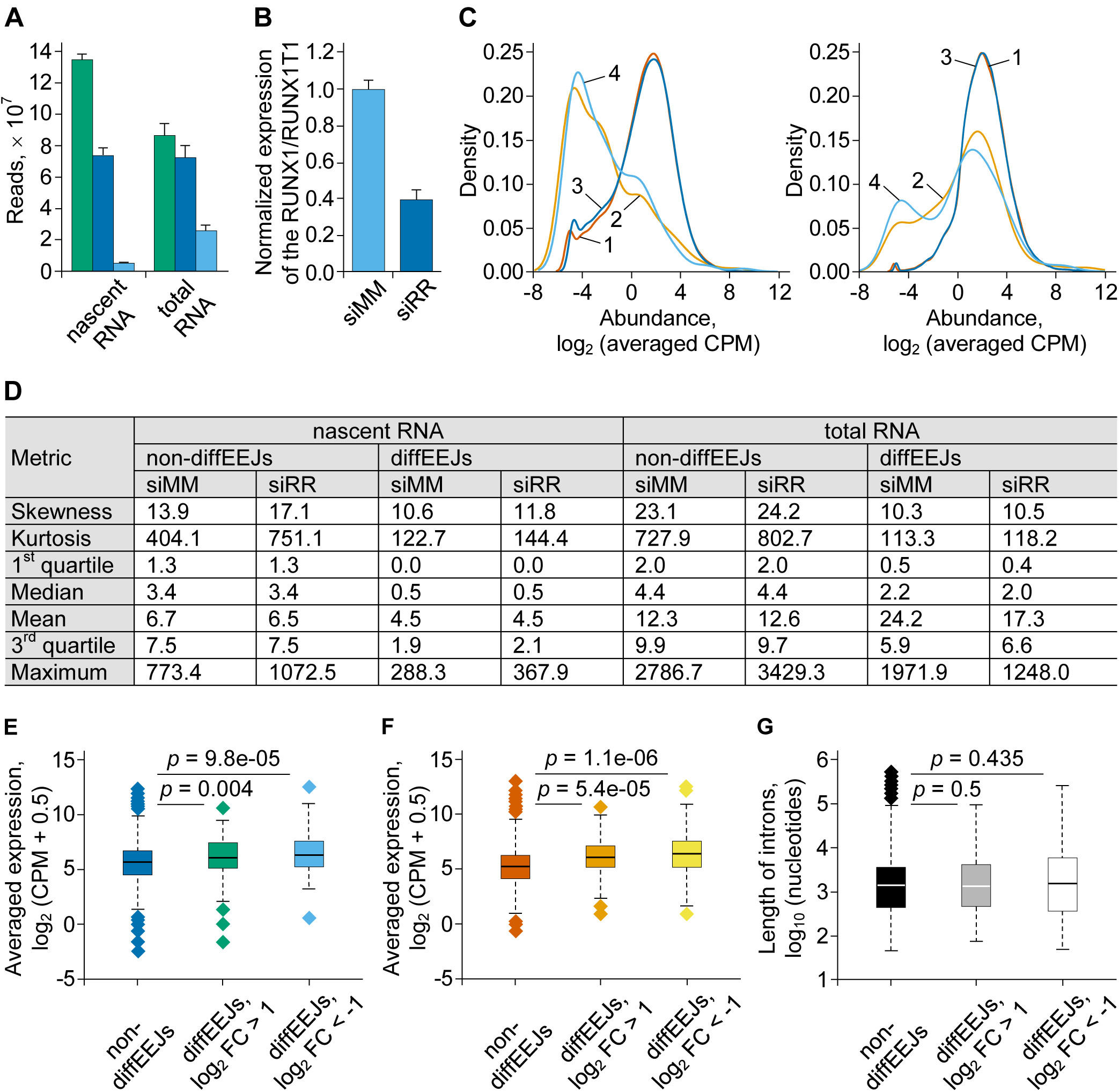
At the nascent RNA level, the formation of diffEEJs is delayed compared to non-diffEEJs. (A) The basic alignment statistics for the nascent RNA and total RNA datasets obtained from the siMM-treated Kasumi-1 cells. This diagram shows the total number of reads per RNA-Seq library (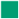), the number of uniquely mapped reads (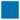), and the number of reads mapped to EEJs only (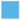). Results are presented as the arithmetic mean plus/minus standard deviation of three independent experiments. The similar results were observed with the RNA datasets obtained from the siRR-treated Kasumi-1 cells. (B) Treatment of Kasumi-1 cells with siRR leads to downregulation of the *RUNX1/RUNX1T1* expression at nascent RNA level. This plot shows the results of qPCR based measuring the expression of the *RUNX1/RUNX1T1* from three independent experiments (as the arithmetic mean plus/minus standard deviation). (C) Multidensity plots of the abundance of EEJs identified in the nascent RNA (left panel) and total RNA (right panel). These plots are based on the RNA-Seq data obtained from the siMM-(lines 1 and 2) or siRR-treated (lines 3 and 4) Kasumi-1 cells. All EEJs identified at the level of nascent RNA and total RNA were grouped (see main text for further explanation of classification procedure) into non-diffEEJs (lines 1 and 3) or diffEEJs (lines 2 and 4), and the abundance of these events was plotted. Each line represents the data averaged over the three independent biological replicas. (D) Basic descriptive statistics on the logarithmically untransformed data from (C). (E) and (F) Genes with differential EEJs tend to higher expression at nascent (E) and total (F) (F) RNA level than genes without EEJs. Boxplots summarize the expression data averaged over the three independent biological replicas. (G) There are no differences in the length of introns between non-diffEEJs and diffEEJs.

Interestingly, expression of diffEEJs (affected by RUNX1/RUNX1T1 knockdown) showed a much stronger bimodal distribution resulting in a lower median compared to non-diffEEJs. For the nascent RNA, the number of low abundant diffEEJs substantially exceeded that of high abundant ones, while the opposite was true for total RNA. In contrast, the distribution pattern of non-EEJs shifted only slightly towards higher abundances (Figure 4C, D). Importantly, neither expression levels of differentially spliced versus non-differentially spliced genes nor the distribution of intron lengths differed between non-diffEEJs and diffEEJs and are, thus, not the cause for the observed shifts (Figures 4E, 4F, 4G). Therefore, these findings suggest the presence of differential splicing events which undergo delayed processing. Furthermore, they imply that delayed splicing is more sensitive to genomic and transcriptomic perturbations induced by the changes in the *RUNX1/RUNX1T1* expression.

### Differential splicing can arise from alternative promoter usage

RUNX1/RUNX1T1 binding sites are enriched in promoter and intronic regions indicating that the fusion protein directly affects the promoter function of multiple genes (Maiques-Diaz et al., 2012; Ptasinska et al., 2012, 2014). In our dataset, we found that 38% of genes with differential splicing are also enriched for the fusion protein binding sites (data not shown). This enrichment is significantly higher than in a sub-set of genes with non-differential splicing (odds ratio 2.47 and p-value = 2.5 × 10^−15^ according to two-sided Fisher’s exact test).

We, thus, hypothesized that at least a part of the identified differential splicing events may arise from the selection of alternative transcription start sites (TSSs) under the control of the fusion protein. Indeed, in the genome of Kasumi-1 cells, we identified 775 differentially expressed TSSs (more than 2-fold change, p < 0.005, q-value < 0.1) following *RUNX1/RUNX1T1* knockdown (Figure 5A). Moreover, we also observed that knockdown of the *RUNX1/RUNX1T1* leads to differential usage of 112 alternative TSSs (more than 2-fold change, p < 0.0007, q-value < 0.1) in the genome of Kasumi-1 cells (Figure 5B). Herewith, the combined analysis of ChIP-Seq and RNA-Seq of total and nascent RNA indicate a direct link between RUNX1/RUNX1T1 binding and the activation of alternative TSSs (Figure 5C).

**Figure 5.**
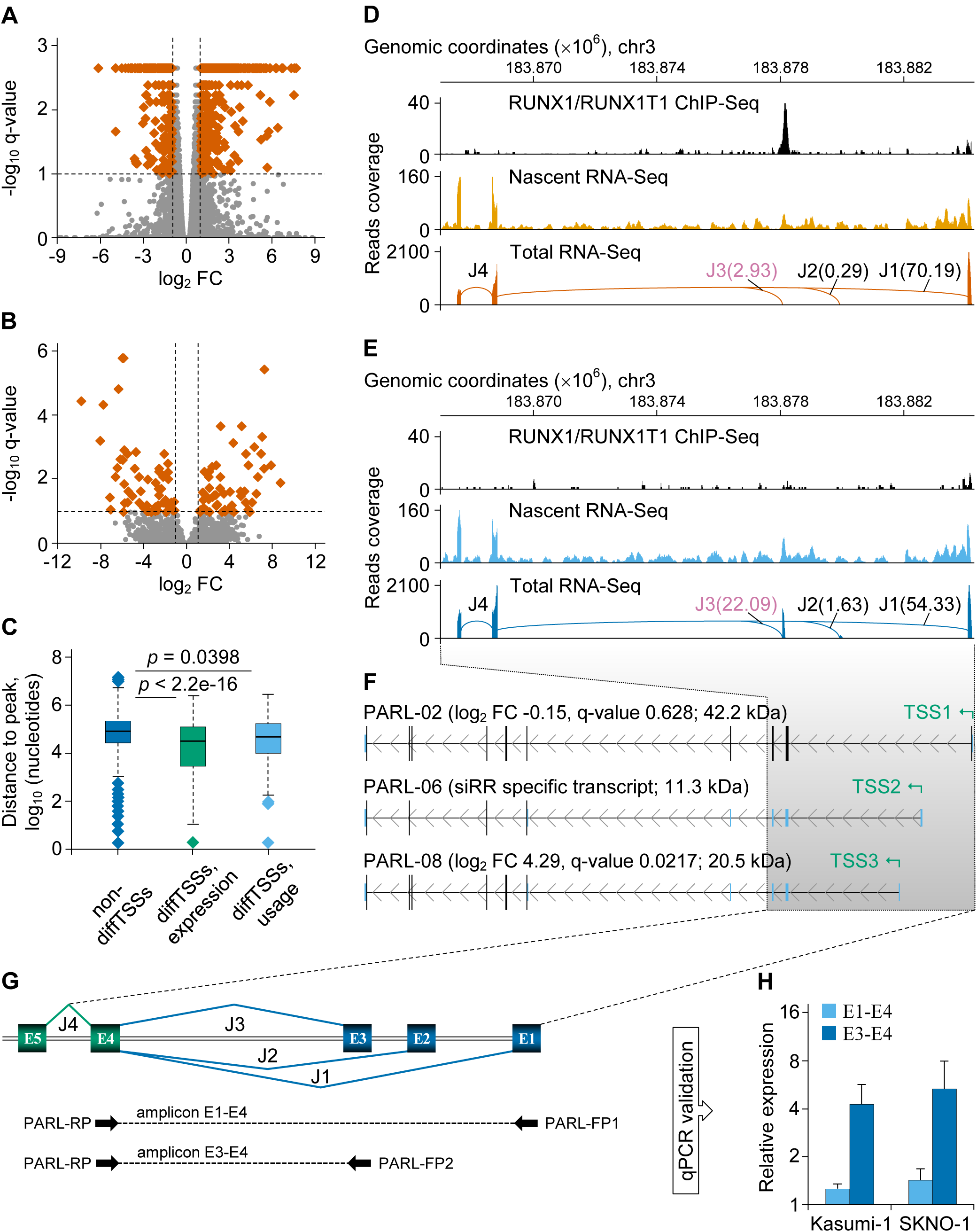
The cis-effect of the fusion protein RUNX1/RUNX1T1 on the differential splicing of *PARL* transcripts. (A) Volcano plot of the differential expression from various TSSs in the genome of Kasumi-1 cells following *RUNX1/RUNX1T1* knockdown. Differentially expressed TSSs (at least 2-fold change in expression, p-value < 0.005, q-value < 0.1) are shown by vermilion squares. (B) Differential usage of alternative TSSs in the genome of Kasumi-1 cells after *RUNX1/RUNX1T1* knockdown. Differentially used TSSs (at least 2-fold change in expression, p-value < 0.0007, q-value < 0.1) are shown by vermilion squares. (C) Statistical summary on proximity of non-differential TSSs (non-diffTSSs), differentially expressed TSSs (diffTSSs, expression) or differentially used TSSs (diffTSSs, usage) to the nearest RUNX1/RUNX1T1 binding peaks in Kasumi-1 cells. The statistical significance of the observed differences was assessed using the two-sided Mann-Whitney U test. (D) In siMM-treated Kasumi-1 cells, RUNX1/RUNX1T1 binds the genomic region near the alternative internal promoter of *PARL* gene and suppresses the transcriptional activity of this promoter. As a result, the frequency of the junction J3 is low in the transcriptome of the control leukemia cells at the level of nascent RNA, as well as in total RNA. Hereinafter in Figure 5E, the splicing graphs of *PARL* gene were reconstructed from the total RNA-Seq reads mapped to exons and EEJs of *PARL* gene. EEJs are designated by the letter J and numbered. The normalized number of reads supporting junctions associated with alternative first exons is shown in parentheses, and the differential splicing event is highlighted in purple. (E) Depletion of the fusion protein leads to augmentation of the internal promoter activity of *PARL* gene in siRR-treated Kasumi-1 cells. As a result, the frequency of the junction J3 increases in the transcriptome of the leukemia cells both at the level of nascent RNA and at the level of total RNA. (F) Cufflinks assembled representative full-length transcripts of *PARL* gene expressed in Kasumi-1 cells. For each transcript, Cuffdiff-based log_2_ FC and q-values as well as the size of *in silico* predicted proteins are indicated in parentheses. In addition, the positions of the Cuffdiff-determined transcription start sites are also shown. (G) Schematic representation of the 5„-terminal region of *PARL* gene. Exons and EEJs are designated by the letters E and J, respectively, and numbered. Binding positions of the qPCR primers and the structure of the expected amplicons are shown at the bottom of the figure. (H) qPCR-based validation of the differential splicing of *PARL* transcripts under the two siRNA treatment conditions and in the transcriptome of two t(8;21)-positive cell lines.

For instance, *PARL* is a direct RUNX1/RUNX1T1 target gene that encodes a mitochondrial protease (Jeyaraju et al., 2011). Knockdown of RUNX1/RUNX1T1 eliminated its binding to an element 6.6 kb downstream of the canonical *PARL* TSS1. This loss of binding activated a 3’-proximally located TSS2 and TSS3. The expression of this novel exon and the corresponding exon-exon junction J3 yielded transcripts PARL-006 and PARL-008 (Figures 5D-F). Interestingly, the observed increase in total *PARL* transcript levels upon RUNX1/RUNX1T1 knockdown was mainly due to these two alternative transcripts (Figures 5G-H). Furthermore, junction J3 is a specific feature of the transcriptomes of at least some primary inv(16) and normal karyotype AMLs, but neither of t(8;21)-positive AMLs nor of normal cord blood (CB-SC) or bone marrow (BM-SC) derived CD34-positive hematopoietic stem/progenitor cells (Figure S3). Additional examples for activation of novel TSSs upon loss of RUNX1/RUNX1T1 binding include *RPS6KA1* and *NADK* (Figure S4). These findings demonstrate the ability of RUNX1/RUNX1T1 to control gene expression by preventing alternative initiation of transcription. They may also provide an explanation for potential discordances between changes in transcript and protein levels following *RUNX1/RUNX1T1* knockdown.

### Differential expression of splicing factors is an additional source of differential splicing in Kasumi-1 cells

Around 60% of differential splicing events identified in Kasumi-1 cells cannot be explained by proximal RUNX1/RUNX1T1 binding indicating that they are not under direct control of the fusion protein. For instance, we identified differential exon-exon junction J5 from the middle part of *TRAPPC2L* gene (Figure 6), which encodes a protein involved in vesicle transport (Barrowman et al., 2010; Scrivens et al., 2009) and which is not a direct target of RUNX1/RUNX1T1 (data not shown). In fact, this region of gene is under a complex and cell type-specific regulation including alternative termination of transcription, alternative transcription start sites, selection of alternative 5’ and/or 3’ splice sites, and cassette exons (Figure S5).

**Figure 6.**
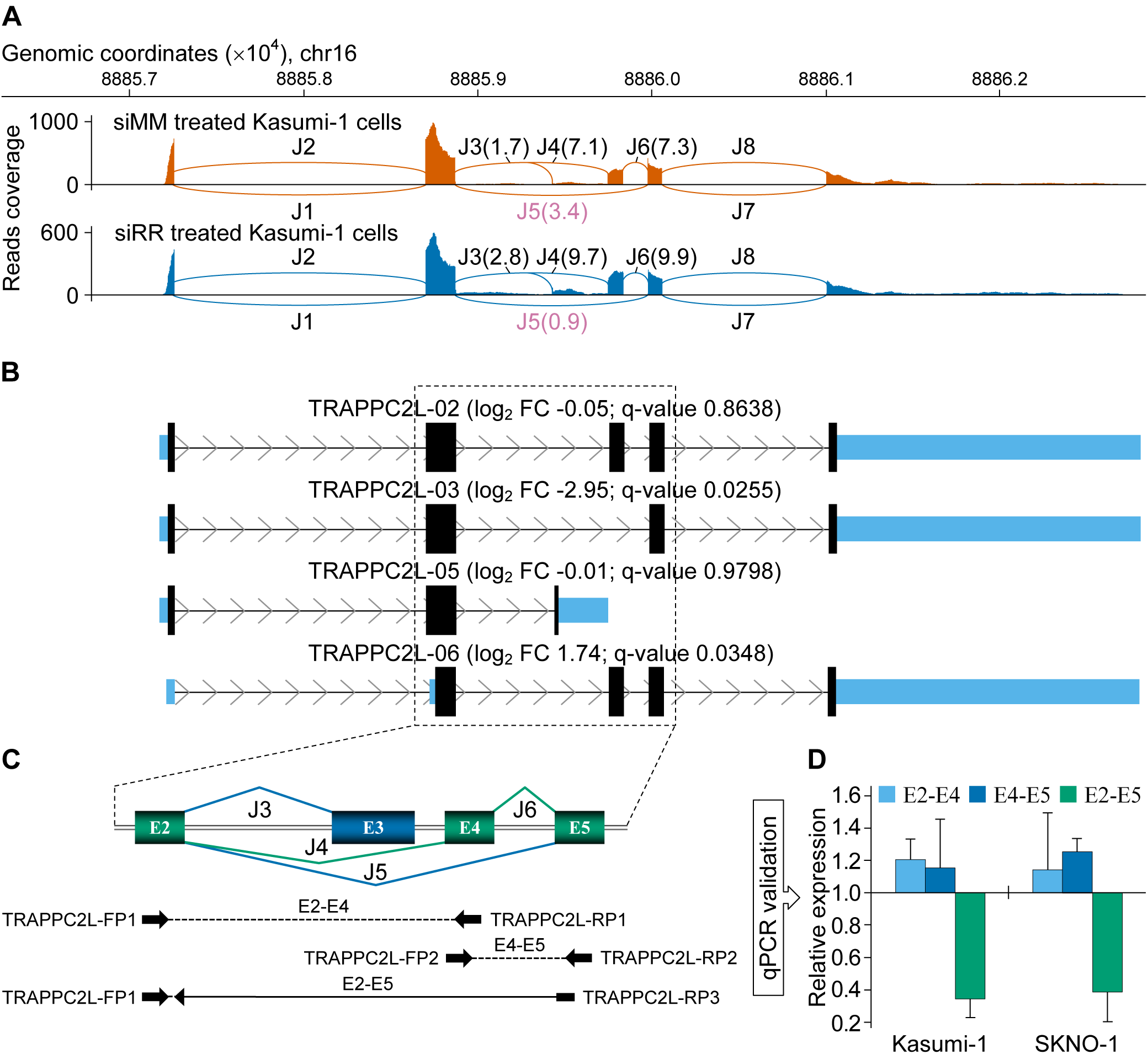
The knockdown of the fusion oncogene *RUNX1/RUNX1T1* leads to differential splicing of TRAPPC2L transcripts. (A) Splicing graphs of *TRAPPC2L* gene. These graphs were reconstructed from RNA-Seq reads mapped to exons and EEJs of *TRAPPC2L* gene. EEJs are designated by the letter J and numbered. The normalized number of reads supporting boxed junctions is shown in parentheses, and the differential splice event is highlighted in purple. (B) Cufflinks assembled representative full-length transcripts of *TRAPPC2L* gene expressed in Kasumi-1 cells. For each transcript, Cuffdiff-based log_2_ FC and q-values are indicated in parentheses. (C) Schematic representation of the target region of *TRAPPC2L* gene. Exons and EEJs are designated by the letters E and J, respectively, and numbered. Binding positions of the qPCR primers and the structure of the expected amplicons are shown at the bottom of the figure. (D) qPCR-based validation of the differential splicing of TRAPPC2L transcripts under the two siRNA treatment conditions and in the transcriptome of two t(8;21)-positive cell lines.

Changed expression of genes encoding splicing factors can change splicing patterns of RNA molecules (Lee et al., 2015; Ramanouskaya et al., 2017). Furthermore, expression of mRNA surveillance genes may also affect the abundance of RNA isoforms (Ge et al., 2014; Weischenfeldt et al., 2012; Yan et al., 2015). Interestingly, we and others have previously shown that t(8;21)-positive leukemia cells have a specific signature of genes encoding splicing factors and mRNA surveillance genes expression (Grinev et al., 2015; Maciejewski et al., 2012). This signature is so unique that it allows distinguishing leukaemic from normal hematopoietic cells (Figures S6A-S6B). Moreover, correlation analysis confirmed highly coordinated expression of genes encoding splicing factors and mRNA surveillance genes with *RUNX1/RUNX1T1* in t(8;21)-harboring AML cells (Figure S6C). Therefore, we speculated that RUNX1/RUNX1T1-dependent changes in expression of RNA processing genes may be responsible for additional differential splicing in Kasumi-1 cells.

By combining of different approaches, we identified 42 genes encoding splicing factors and 14 mRNA surveillance genes with statistically significant differential expression in Kasumi-1 cells following the *RUNX1/RUNX1T1* knockdown (Table S7, Table S8, Figure 7A). Many of those are associated with RUNX1/RUNX1T1 binding sites implying them as direct target genes. The *USB1* gene is an example for this group of genes (Figure 7B). Nevertheless, genes encoding splicing factors and mRNA surveillance genes may also indirectly be regulated via fusion protein-mediated control of transcription factors expression.

**Figure 7.**
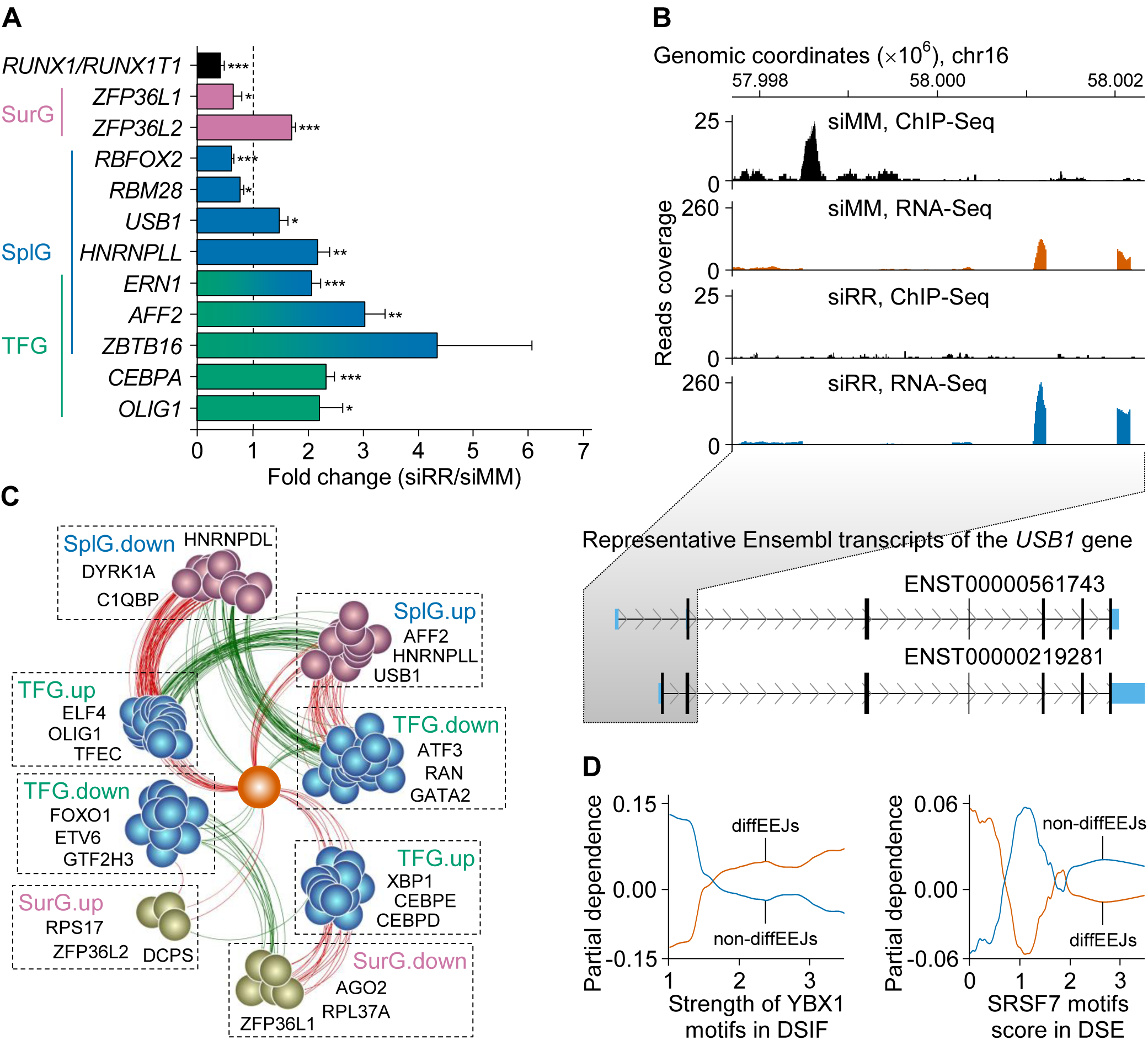
The indirect effect of the fusion protein RUNX1/RUNX1T1 on the differential splicing in Kasumi-1 cells. (A) The knockdown of the fusion oncogene RUNX1/RUNX1T1 leads to differential expression of mRNA surveillance genes (SurG), genes encoding splicing factors (SplG) and genes of transcription regulators (TFG). According to Gene Ontology, three of these genes (*AFF2*, *ERN1* and *ZBTB16*) demonstrate dual transcriptional and splicing regulatory activity. Each bar represents the arithmetic mean plus/minus standard deviation of three independent qPCR experiments. Student t-test was used to check out of statistical significance: (*) p < 0.05, (**) p < 0.01, (***) p < 0.001. (B) *USB1* gene is a direct target of RUNX1/RUNX1T1. According to the ChIP-Seq and RNA-Seq analysis, fusion protein binds 459 nucleotides upstream of the first transcription start site and negatively controls the expression of *USB1* gene. (C) RUNX1/RUNX1T1-centered (vermillion ball) gene regulatory network of genes encoding splicing regulators and mRNA surveillance factors. In this plot, different groups of genes are designated by the same abbreviations as in part A of the figure. Terms “up(-regulated)” and “down(-regulated)” refer to a change in the expression of the genes after the *RUNX1*/*RUNX1T1* knockdown. Positively and negatively co-expressed genes linked by green and red edges, respectively. Some examples of the genes from each group are shown in the respective dashed boxes. Graphical view of this network was obtained with Cytoscape. (D) Effect of the strength of the motifs to splicing factors on the probability of non-diffEEJs and diffEEJs in Kasumi-1 cells. These partial dependence plots are based on the results from 1000 independent runs of the random forest meta-classifier and 1000 classification trees per random forest per run.

To check this hypothesis, we reconstructed gene regulatory network for those genes encoding splicing factors and mRNA surveillance genes with significantly changed expression upon RUNX1/RUNX1T1 depletion (Figure 7C). This network was inferred from microarray, RNA-Seq and bioinformatics data (see Supplemental Experimental Procedures for further details). The reconstructed network strongly suggest complex expression interplay between the fusion protein, transcription factors and the factors involved in splicing and mRNA surveillance. For example, the promoter of *HNRNPLL*, whose encoded protein is involved in alternative splicing, is significantly enriched with the motif for the OLIG1 transcription factor (enrichment score 2.8, p-value = 0.0329), and there is strong positive co-expression between the *HNRNPLL* and *OLIG1* genes (robust correlation coefficient is 0.967). In turn, the *OLIG1* gene itself is under direct negative control of RUNX1/RUNX1T1. Consequently, the expression of *HNRNPLL* gene is more than twofold increased (q-value < 0.001) in siRR-treated Kasumi-1 cells.

Based on the above-mentioned predictive model, we developed a short list of sequence features that represent the most important predictors of whether an exon-exon junction is differential or not. This short list included motif frequency and strength for HNRNPA3, HNRNPF, HNRNPH1, MBNL1, PTBP1, SRSF7, TARDBP, TIAL1, TRA2B and YBX1 splicing factors (Table S9). Partial dependence profiling indicated a strong non-linear relationship between class probability of EEJs and frequency or strength of splicing regulator motifs. For example, there is a positive link between the motif strength of the YBX1 splicing factor in the 3’-end of the intron and the probability of diffEEJs, and a negative relationship between SRSF7 motif score in the downstream exon and differential splicing (Figure 7D). Motifs for all the other splicing factors from the above list similarly affected the discrimination between differential versus non-differential splicing events (data not shown).

The splicing factors can be classified according to their response to RUNX1/RUNX1T1 status. One group comprises several hnRNPs (HNRNPDL, HNRNPLL and HNRNPR) with changed expression levels upon RUNX1/RUNX1T1 knockdown, while other hnRNPs such as HNRNPA3, HNRNPF and HNRNPH1 were not affected (Table S7). Interestingly, the SRSF7 motif was also identified as an important feature, but the expression of the respective gene is unchanged in Kasumi-1 cells.

To assess the involvement of RNA surveillance factors in differential splicing, we compared the distribution of RNA isoform expression with and without premature stop codons for all the genes with differential splicing. This analysis revealed no significant difference between the wild type and knockdown conditions (data not shown).

Taken together, these findings suggest that RUNX1/RUNX1T1 dysregulates the expression of splicing factors thereby impacting on differential splicing. Since this association requires RUNX1/RUNX1T1-mediated differential expression of genes encoding splicing factors, rather than the interaction of fusion protein with the alternatively spliced gene, this activity constitutes an indirect effect of RUNX1/RUNX1T1 on differential splicing.

### Differential splicing produces proteins with unique conserved domain structures and affects multiple cellular processes

Finally, we assessed the potential functional consequences of RUNX1/RUNX1T1-associated differential splicing. Enrichment analysis with Gene Ontology gene sets suggested that differential splicing particularly affects genes controlling synthesis, transport, targeting and localization of proteins, cell adhesion and granulocyte activation (Figure 8A).

**Figure 8.**
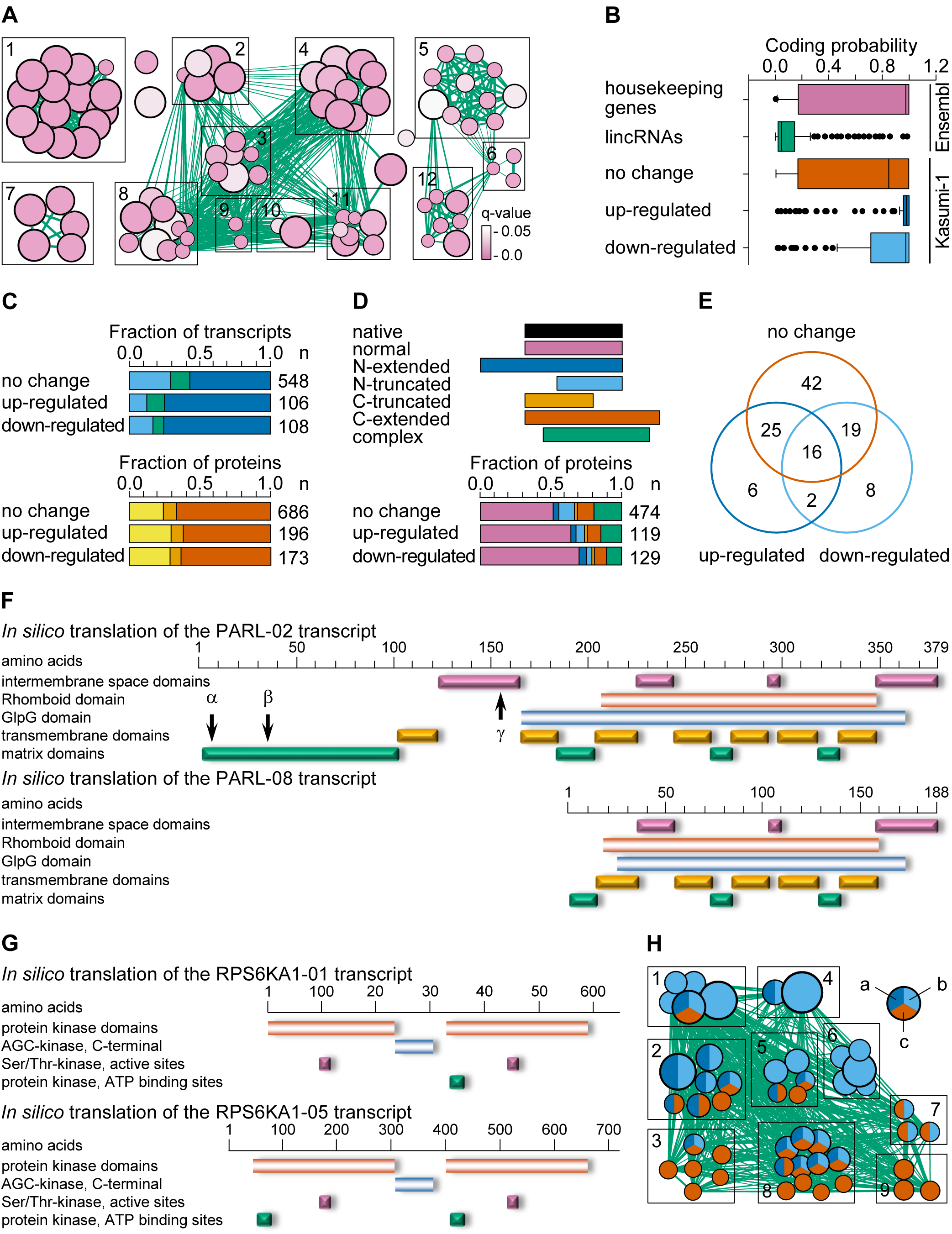
The differential usage of EEJs leads to the appearance of RNA isoforms which encode proteins with unique conserved domain structures in Kasumi-1 cells. (A) Enrichment map of the Gene Ontology-enriched gene sets across all the genes with differential expression and/or differential splicing. Nodes represent significantly enriched gene sets and node size is proportional to the number of members in a gene set. Edges indicate the gene overlap between the nodes, and the thickness of the edges is equivalent to the degree of the gene overlap between the nodes. Functionally related gene sets are clustered and named: 1) granulocyte activation response, 2) virus-cell interaction, 3) mRNA/rRNA metabolic process, 4) amide metabolic process, 5) purine metabolic process, 6) carbohydrate catabolic process, 7) cell-substrate adherents, 8) protein targeting and localization, 9) co-translation protein targeting, 10) intracellular protein transport, 11) protein synthesis, 12) pyruvate metabolic process. (B) The coding potential of 500 randomly selected transcripts of the housekeeping genes, 500 randomly selected lincRNAs, and all transcripts of the genes affected by differential splicing in Kasumi-1 cells. (C) Identification and classification of coding sequences in transcripts of the genes affected by differential splicing in Kasumi-1 cells. In the upper panel, all transcripts were classified on non-coding (no significant open reading frames were found, 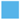), protein-coding with premature termination codons (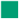) and protein-coding with mature termination codons (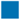). In the bottom panel, all protein-coding transcripts were *in silico* translated and predicted proteins were blastp aligned against non-redundant set of human proteins. According to alignment results, all the *in silico* predicted proteins were classified on no-hits (no significant identities with canonical proteins were found, 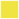), lowly identical (with identity below 90%, 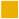) and highly identical (identity above 90%, 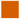). (D) A substantial fraction of highly identical proteins is represented by native human proteins. At the same time, there are different extended and truncated variants of proteins. (E) Distribution of the 118 domain superfamilies among the *in silico* predicted proteins. (F) Domain architecture of the *in silico* translated two PARL proteins. This picture is compiled from the outputs of NCBI Conserved Domains and EMBL-EBI InterPro services. The cleavage sites of the full-length isoform of the PARL protein are marked by arrows. (G) Domain architecture of the *in silico* translated two RPS6KA1 proteins. This picture is compiled from the outputs of NCBI Conserved Domains and EMBL-EBI InterPro services. (H) Enrichment map of the domain-centered Gene Ontology-enriched superfamily sets across all the genes with differential splicing. Nodes represent significantly enriched superfamily sets and node size is proportional to the number of members in a superfamily set. Edges indicate the superfamily overlap between the nodes, and the thickness of the edges is equivalent to the degree of the superfamily overlap between the nodes. Functionally related superfamily sets are clustered and named: 1) regulation of cellular physiological process, 2) regulation of biological process, including enzyme activity, 3) regulation of response to stress, including immune response, 4) molecular binding, including protein binding, 5) molecular transport and localization of molecular substances, 6) plasma membrane and cell periphery, 7) regulation of cell communication, 8) regulation of cell differentiation and cellular development process, 9) signal transduction. Pie chart coloring: (a) up-regulated, (b) down-regulated, and (c) no change.

To further detail above functional findings, we used the Cufflinks-assembled transcriptome of Kasumi-1 cells to select all 762 transcripts of the genes affected by RUNX1/RUNX1T1-dependent differential splicing. These genes gave rise to 214 differentially spliced and 548 non-differentially spliced RNA isoforms. Here and below, the term “differentially spliced isoform” indicates a molecule of RNA with at least one differential exon-exon junction (up- or down-regulated), while a non-differential isoform is a transcript lacked any diffEEJs.

We found that 75% of the all above transcripts were potentially protein coding, while the rest were non-coding RNAs (Figure 8B; Figure 8C, upper panel). We aligned *in silico* translated proteins against the human up-to-date NCBI RefSeq proteins and the newest non-redundant releases of GenBank CDS translations, UniProtKB/SwissProt, Protein Data Bank, Protein Information Resource and Peptide/Protein Sequence Database proteins using NCBI blastp (Gish et al., 1993). Out of 1055 *in silico* translated proteins, 722 proteins were 90% or more identical to the proteins annotated in the public databases (Figure 8C, bottom panel). Such proteins form a clear peak on the curve of identity distribution (data not shown) and, to reduce noise, only these proteins were selected for downstream analysis.

A substantial fraction of *in silico* translated proteins is represented by native human proteins (Figure 8D). At the same time, there are different extended and truncated isoforms of proteins. Such isoforms can potentially include new domains or can be lacked domains important for proper protein binding, localization and function, whereas other functional regions remained unaffected.

To further structural detail, we assigned SCOP domains for highly identical *in silico* translated proteins using the SUPERFAMILY hidden Markov models, and we identified 225 conserved domains within these proteins (Structural Classification of Proteins database, release 1.75, June 2009; Hubbard et al., 1999; Wilson et al., 2009). Identified domains belong to 146 domain families. About 45% of these domain families are common for proteins translated from differentially (up- or down-regulated) and non-differentially (no change) spliced RNA isoforms. However, 16% of domain families are associated with differentially spliced RNA isoforms. This uniqueness of differential proteins is robust and is retained even when domains are combined into domain superfamilies (Figure 8E).

The *PARL* gene provides a good example of a gene producing multiple proteins with different domain architectures. For instance, transcript PARL-02 is non-differentially spliced and non-differentially expressed in Kasumi-1 cells. This transcript encodes full-length 42 kDa protein, which is identical to NCBI RefSeq protein NP_061092.3. At the same time, transcript PARL-08, which is a product of a differential splicing event, potentially encodes an N-terminally truncated 20 kDa isoform of PARL protein. This predicted isoform is completely devoid of the main cytoplasmic domain and the first transmembrane and extracellular domains. Furthermore, it would contain a truncated second cytoplasmic domain and truncated conserved membrane-associated serine protease domain GlpG from rhomboid superfamily (Figure 8F). The *RPS6KA1* gene is another example of genes encoding full-length and N-truncated proteins (Figure 8G). However, in this case the differentially expressed transcript RPS6KA1-05 encodes a full-length protein containing both N-terminal the intact protein kinase domain and the ATP binding site.

Finally, we used a domain-centric Gene Ontology method (De Lima Morais et al., 2011) for functional annotation of the domain architectures of *in silico* predicted proteins. We applied the enrichment test separately for proteins translated from differentially spliced (up- or down-regulated) or non-differentially spliced RNA isoforms, and we integrated the results using EnrichmentMap (Merico et al., 2010). This approach showed that the domains encoded by differential RNA isoforms are mainly involved in the global regulation of cellular processes, molecular binding (including protein binding), transport and localization of molecular substances, and cell communications (Figure 7H).

Thus, RUNX1/RUNX1T1 directly and indirectly regulates alternative RNA splicing for a sub-set of genes in the t(8;21)-positive AML cells. This RUNX1/RUNX1T1-mediated differential splicing affects several functional groups of genes and may produce proteins with unique conserved domain structures and functional activity.

## DISCUSSION

Previous *RUNX1/RUNX1T1*-centered transcriptomic studies focused on detecting gene effects, in which entire genes are up- or downregulated depending on the expression of the fusion oncogene. Here, we have extended these studies by analysing the impact of RUNX1/RUNX1T1 on the generation of alternative transcripts by controlling promoter choice and affecting splicing.

Identification of differential splicing events using RNA-Seq data is not a trivial task and there is no universal and perfectly reliable bioinformatic tool for this (Anders et al., 2012; Hartley et al., 2016; Shen et al., 2012). For this reason, we combined several algorithms to detect a comprehensive set of differential splicing events of various modes at exon or exon-exon junction levels with subsequent experimental validations. Using this approach, we identified 378 genes with significantly different RNA splicing maps in Kasumi-1 cells following the fusion oncogene knockdown. A part of the identified differential splicing events arises from the differential usage of alternative transcription starts and is mediated by the fusion protein binding. This agrees with the observation that transcription factors regulate alternative splicing by occupying and modulating alternative promoters (Davuluri et al., 2008; Jacox et al., 2010; Ma et al., 2009).

It was recently shown that RUNX1/RUNX1T1 may recruit splicing regulatory proteins including proteins with poly(A) RNA binding activity (i.e. CIRBP, DKC1, GAR1 or RBM3) (Mandoli et al., 2016). These data suggest that the fusion protein may potentially control polyadenylation. Indeed, we found 614 3’-terminal exons that overlap with RUNX1/RUNX1T1 binding peaks (data not shown). However, only 4 of these exons were differentially used in the transcriptome of the siRR-versus siMM-treated Kasumi-1 cells (at least 2 fold changes, q-value < 0.0001). Thus, RUNX1/RUNX1T1 does not substantially affect the selection of mRNA 3’-termini.

At the same time, the fusion protein may recruit classical splicing regulatory proteins, especially from the SR (SRSF3 and SRSF7) and RBM (RBM3, RBM28, RBM42, RBMS1 and RBMX) families (Mandoli et al., 2016). This could directly lead to differential splicing in the transcriptionally active regions which overlap with the binding peaks for the fusion protein. The SRSF7 motif is an important predictor of RUNX1/RUNX1T1-dependent differential splicing thus supporting this notion. However, the expression of the *SRSF7* gene itself is not altered after the *RUNX1/RUNX1T1* knockdown. We speculate that a change in *RUNX1/RUNX1T1* expression affects the SRSF7 recruitment to splicing area and leads to differential splicing.

Additionally, we found a sub-set of splicing-related genes that were differentially expressed upon RUNX1/RUNX1T1 knockdown. This sub-set includes genes encoding classical splicing regulators (for example, HNRNPLL) as well as multifunctional genes (for instance, DYRK1A) encoding proteins with different activities, including components of the nuclear speckles, which are considered an area of intensive splicing (Cardoso et al., 2012; Lamond et al., 2003). According to our data mining and motif analysis results, these splicing regulatory proteins may additionally contribute to differential splicing in Kasumi-1 cells following the *RUNX1/RUNX1T1* knockdown.

In conclusion, we have shown here that RUNX1/RUNX1T1 affects alternative splicing of RNA in the t(8;21)-positive AML cells by controlling the choice of transcriptional start sites and by modulating expression of splicing components. This activity of the fusion oncogene affects RNA and protein isoforms expressed in the leukemia cells and, thereby, domain architecture of the proteins, thus adding another layer of dysregulatory function promoting leukaemogenesis.

## EXPERIMENTAL PROCEDURES

More detailed descriptions of the materials and methods used in this work can be found in the Supplemental Experimental Procedures.

### Cell lines

The t(8;21)-positive AML cell lines Kasumi-1 (DSMZ no. ACC 220) and SKNO-1 (DSMZ no. ACC 690) were obtained from the DSMZ (LGC Standards GmbH, Wesel, Germany) and cultivated according to the standard approaches.

### siRNA transfections

Kasumi-1 and SKNO-1 cells were transfected with 200 nM siRNA using a Fischer EPI 3500 electroporator (Fischer, Heidelberg, Germany) as described previously (Martinez et al., 2004). In all the RUNX1/RUNX1T1 knockdown experiments, the anti-RUNX1/RUNX1T1 active siRR (sense strand 5’-CCUCGAAAUCGUACUGAGAAG-3’ and antisense strand 5’-UCUCAGUACGAUUUCGAGGUU-3’) and mismatch (inactive) control siMM (sense strand 5’-CCUCGAAUUCGUUCUGAGAAG-3’ and antisense strand 5’-UCUCAGAACGAAUUCGAG GUU-3’) were used (Heidenreich et al., 2003).

### Microarray and high-throughput sequencing

The time series of microarray data on siRR- and siMM-treated Kasumi-1 cells was generated previously (Ptasinska et al., 2012) using Illumina HumanHT-12 V4.0 expression beadchip. Procedure for total RNA isolation, preparation of RNA-Seq libraries, and massively parallel sequencing was previously described in (Ptasinska et al., 2014). Distribution of the DNase I hypersensitivity sites and positions of the H3K9Ac, RNA polymerase II, and RUNX1/RUNX1T1 peaks in the genome of Kasumi-1 cells were previously established and described (Ptasinska et al., 2012).

### Extraction of total cellular RNA, synthesis of the first strand of cDNA and qPCR

Total cellular RNA was extracted using the RNeasy Mini Kit (QIAGEN GmbH, Hilden, Germany) according to the manufacturer’s protocol. Synthesis of the first strand of cDNA was performed from 1 μg of total cellular RNA in 20 μl volume using oligo(dT)18 primer and SuperScript^TM^ III Reverse Transcriptase Kit (Thermo Fisher Scientific, Carlsbad, USA) according to the manufacturer’s protocol. Real-time qPCR was carried out in triplicates on StepOnePlus Real-time PCR System (Thermo Fisher Scientific, Carlsbad, USA) using QuantiTect^®^ SYBR^®^ Green PCR Kit (QIAGEN GmbH, Hilden, Germany) according to the manufacturer’s protocol. All the primers (Table S5) for real-time qPCR were in-laboratory designed using Primer-BLAST on-line tool (Ye et al., 2012).

### Extraction of total cellular proteins and Western blotting

Total cellular proteins were extracted simultaneously with the RNeasy Mini Kit-based purification of RNA using acetone precipitation. Pelleted proteins were dissolved in 9M urea buffer and protein concentration was measured by Bradford assay.

Western blotting was carried out according to the previously described protocol (Heidenreich et al., 2003). Mouse monoclonal anti-PARL IgG_1_ (cat. #sc-514836, Santa Cruz Biotechnology Inc., Santa Cruz, USA), raised against the C-terminus of human PARL, were used as primary antibodies for detection of the PARL protein isoforms. In addition, mouse anti-GAPDH monoclonal antibodies (cat. #5G4 MAb 6C5, HyTest Ltd., Turku, Finland) were used as primary antibodies recognizing GAPDH. Goat anti-mouse polyclonal antibodies conjugated with horseradish peroxidase (cat. #P0447, Agilent, Santa Clara, USA) were used as secondary antibodies.

### Nascent RNA-Seq

Procedure for nascent RNA isolation, preparation of RNA-Seq libraries, and massively parallel sequencing was previously described (Kerry et al., 2017). Briefly, 10^8^ Kasumi-1 cells were treated with 500 μM 4-thiouridine for 1 hour. Cells were lysed using TRI Reagent^®^ (Sigma-Aldrich, St Louis, USA) and RNA was purified according to the manufacturer’s instruction. Next, 4-thiouridine-incorporated RNA was biotinylated by labelling with 1 mg/ml Biotin-HPDP (Abcam, Cambridge, UK) for 90 minutes at room temperature. Following chloroform extraction, labelled RNA was separated using magnetic streptavidin beads, beads were washed and RNA was eluted in two rounds of elution with 100 μl 100 mM DTT. Finally, RNA was purified using RNeasy MinElute Cleanup Kit (QIAGEN GmbH, Hilden, Germany) and samples were sequenced using Illumina HiSeq^TM^ 2000 paired-end sequencing.

### Publicly available high-throughput datasets used in the study

Our public microarray dataset included data on 106 primary bone marrow or peripheral blood samples from the patients with t(8;21)-positive AML, and 85 samples of BM-SC, 32 samples of BM-MNC and 100 samples of PB-MNC from healthy donors. Our public RNA-Seq dataset included data on primary samples from 20 patients with t(8;21)-positive AML, 2 patients with cytogenetically normal AML, 2 patients with inv(16)-positive AML, 2 samples of normal CB-SC and 2 samples of normal BM-SC.

### Annotated public datasets

All annotations of human genome were downloaded from Ensembl database (Zerbino et al., 2018) in GTF/GFF format. We used Ensembl release 85 based on GRCh38.p7 reference assembly of human genome. Genomic coordinates of the CpGs islands in the human genome were downloaded via FTP server of UCSC Genome Browser (Speir et al., 2016). The whole set of GenBank human mRNA and ESTs sequences was downloaded via FTP server of UCSC Genome Browser in GTF format.

### Analytical approaches

All our analytical approaches are described in details in the Supplemental Experimental Procedures. Our analytical pipeline was mainly based on the capabilities of the R programming language and Bioconductor facilities.

## Supporting information

Supplemental Experimental Procedures

## ACCESSION NUMBERS

The accession numbers for all the microarray and next generation sequencing data reported in this paper is listed in Table S1.

## SUPPLEMENTAL INFORMATION

Supplemental Information includes Supplemental Experimental Procedures, seven figures, and seven tables and can be found with this article online.

## AUTHOR CONTRIBUTIONS

V.G., I.I., R.G. and N.M.-S. performed experiments. V.G., I.I. and S.N. analyzed data. T.R. contributed to preparation and editing of the manuscript. C.B. and O.H. conceived the study, supervised experiments, and wrote the manuscript.

## ACKNOWLEDGEMENTS

The authors thank Aliaksandra Radzisheuskaya (Biotech Research and Innovation Centre, University of Copenhagen, Copenhagen, Denmark) for carefully reading and improving the manuscript. Research in the V.G. laboratory was supported in part by the Ministry of Education of the Republic of Belarus, grant #3.08.3.

