## Supplemental Experimental Procedures for "RUNX1/RUNX1T1 controls alternative splicing in the t(8;21)-positive acute myeloid leukemia cells"

### 1. Cell and molecular biology methods

#### 1.1. Cell lines

The t(8;21)-positive AML cell lines Kasumi-1 (DSMZ no. ACC 220) and SKNO-1 (DSMZ no. ACC 690) were obtained from the DSMZ (LGC Standards GmbH, Wesel, Germany). Leukemia cells were cultivated in the RPMI 1640 medium with the addition of 10% fetal bovine serum, 2 mM L-glutamine, 100 U/ml penicillin, 100 µg/ml streptomycin and 10 ng/ml GM-CSF (for SKNO-1 cells) at 37°C in the atmosphere with 5% $CO_2$.

#### 1.2. siRNA transfections

Kasumi-1 and SKNO-1 cells were transfected with 200 nM siRNA using a Fischer EPI 2500 electroporator (Fischer, Heidelberg, Germany) as described previously (Martinez et al., 2004). In all the *RUNX1*/*RUNX1T1* knockdown experiments, the anti-RUNX1/RUNX1T1 active siRR (sense strand 5'-CCUCGAAAUCGUACUGAGAAG-3' and antisense strand 5'-UCUCAGUACGAUUUCGAGGUU-3') and mismatch (inactive) control siMM (sense strand 5'-CCUCGAAUUCGUUCUGAGAAG-3' and antisense strand 5'-UCUCAGAACGAAUUCGAGGUU-3') were used (Heidenreich et al., 2003).

#### 1.3. Extraction of total cellular RNA

Total cellular RNA was extracted using the RNeasy Mini Kit (QIAGEN GmbH, Hilden, Germany) according to the manufacturer's protocol. RNA concentration was determined by loading 1 µl of sample on a NanoDrop$^{TM}$ 1000 Spectrophotometer (Thermo Fisher Scientific, Carlsbad, USA).

#### 1.4. Synthesis of the first strand of cDNA and real-time qPCR

Synthesis of the first strand of cDNA was performed from 1 µg of total cellular RNA in 20 µl volume using oligo(dT)18 primer and SuperScript$^{TM}$ III Reverse Transcriptase Kit (Thermo Fisher Scientific, Carlsbad, USA) according to the manufacturer's protocol. Real-time qPCR was carried out in triplicates of 20 µl volume with 0.3 µM of each primer and 1 µl of total cDNA as a template on StepOnePlus Real-time PCR System (Thermo Fisher Scientific, Carlsbad, USA) using QuantiTect® SYBR® Green PCR Kit (QIAGEN GmbH, Hilden, Germany) according to the manufacturer's protocol. All the primers (Table S5) for real-time qPCR were in-laboratory designed using Primer-BLAST on-line tool (Ye et al., 2012).

#### 1.5. Extraction of total cellular proteins

Total cellular proteins were extracted simultaneously with the RNeasy Mini Kit-based purification of RNA using acetone precipitation. Pelleted proteins were dissolved in 9M urea buffer and protein concentration was measured by Bradford assay.

#### 1.6. Western blotting

Western blotting was carried out according to the previously described protocol (Heidenreich et al., 2003). Mouse monoclonal anti-PARL $IgG_1$ (cat. #sc-514836, Santa Cruz Biotechnology Inc., Santa Cruz, USA), raised against the C-terminus of human PARL, were used as primary antibodies for detection of the PARL protein isoforms. In addition, mouse anti-GAPDH monoclonal antibodies (cat. #5G4 MAb 6C5, HyTest Ltd., Turku, Finland) were used as primary antibodies recognizing GAPDH. Goat anti-mouse polyclonal antibodies conjugated with horseradish peroxidase (cat. #P0447, Agilent, Santa Clara, USA) were used as secondary antibodies.

### 2. Datasets

#### 2.1. In-laboratory generated high throughput datasets

##### 2.1.1. Microarray data

The time series of microarray data on siRR- and siMM-treated Kasumi-1 cells was generated previously (Ptasinska et al., 2012) using Illumina HumanHT-12 V4.0 expression beadchip. Processed data can be downloaded from the NCBI GEO repository (Barrett et al., 2013) under the series accession number GSE29223.

##### 2.1.2. Nascent RNA-Seq data

Procedure for nascent RNA isolation, preparation of RNA-Seq libraries, and massively parallel sequencing was previously described (Kerry et al., 2017). Briefly, $10^8$ Kasumi-1 cells were treated with 500 μM 4-thiouridine for 1 hour. Cells were lysed using TRI Reagent® (Sigma-Aldrich, St Louis, USA) and the RNA was purified according to the manufacturer's instruction. Next, 4-thiouridine-incorporated RNA was biotinylated by labelling with 1 mg/ml Biotin-HPDP (Abcam, Cambridge, UK) for 90 minutes at room temperature. Following chloroform extraction, labelled RNA was separated using magnetic streptavidin beads, beads were washed and RNA was eluted in two rounds of elution with 100 μl 100 mM DTT. Finally, RNA was purified using RNeasy MinElute Cleanup Kit (QIAGEN GmbH, Hilden, Germany) and samples were sequenced using Illumina HiSeq™ 2000 paired-end sequencing.

##### 2.1.3. Total RNA-Seq data

The procedure for total RNA isolation, preparation of RNA-Seq libraries, and massively parallel sequencing was previously described (Ptasinska et al., 2014). FASTQ files with raw data from the siRR- and siMM-treated Kasumi-1 cells are freely available in Sequence Read Archive or European Nucleotide Archive under the study PRJNA236604.

##### 2.1.4. DNase I hypersensitivity sites data

Distribution of the DNase I hypersensitivity sites in the genome of Kasumi-1 cells was previously established (Ptasinska et al., 2012). Raw data are freely available in Sequence Read Archive or European Nucleotide Archive under the study PRJNA142965. Processed data can be downloaded from the NCBI GEO repository (Barrett et al., 2013) as a part of series GSE29222.

##### 2.1.5. ChIP-Seq data

Positions of the H3K9Ac, RNA polymerase II, and RUNX1/RUNX1T1 peaks in the genome of the siRR- and siMM-treated Kasumi-1 cells were previously mapped (Ptasinska et al., 2012). Raw data are freely available in Sequence Read Archive or European Nucleotide Archive under the study PRJNA142965. Processed data can be downloaded from the NCBI GEO repository (Barrett et al., 2013) under the series accession number GSE29222.

#### 2.2. Public high throughput datasets

##### 2.2.1. Microarray data

Our public microarray dataset included data on 106 primary samples of bone marrow or peripheral blood from patients with t(8;21)-positive AML, and 85 samples of BM-SC, 32 samples of BM-MNC and 100 samples of PB-MNC from healthy donors. All the microarray samples are listed and briefly described in Table S1. CEL files of the selected microarrays were downloaded from the NCBI GEO repository (Barrett et al., 2013), converted into CHP files, and primary data matrix was corrected against the background, $log_2$-transformed and quantile normalized by the robust multi-array averaging algorithm using Affymetrix Expression Console™ (Thermo Fisher Scientific, Carlsbad, USA).

##### 2.2.2. Total RNA-Seq data

Our public leukemia dataset included total RNA-Seq data on primary samples from 20 patients with t(8;21)-positive AML, 2 patients with cytogenetically normal AML and 2 patients with inv(16)-positive AML. Moreover, total RNA-Seq data from 2 samples of CB-SC and 2 samples of BM-SC were also used in this study. All these data were generated using Illumina HiSeq™ 2000/2500 paired-end sequencing. Raw data were downloaded from European Nucleotide Archive in a form of FASTQ files. All the total RNA-Seq samples are listed and briefly described in Table S1.

#### 2.3. Annotated public datasets

All the annotations of human genome were downloaded from Ensembl database (Zerbino et al., 2018) in GTF/GFF format. We used Ensembl release 85 based on GRCh38.p7 reference assembly of human genome.

Genomic coordinates of the CpGs islands in the human genome were downloaded via FTP server of the UCSC Genome Browser (Speir et al., 2016).

The whole set of GenBank human mRNA and ESTs sequences was downloaded via FTP server of the UCSC Genome Browser (Speir et al., 2016) in GTF format. We used sequences that were pre-aligned against the GRCh38/hg38 reference assembly of the human genome with BLAT (Kent, 2002). Any sequences with mismatches and less than four aligned blocks were removed. Additionally, sequences with the exon length <25 bp and intron length <50 bp were filtered out.

### 3. Analytical approaches

#### 3.1. Quality assessment and pre-processing of the RNA-Seq raw data

A comprehensive RNA-Seq data quality assessment and pre-processing of the raw data was performed using the R/Bioconductor library ShortRead v.1.38.0 (Morgan et al., 2009) and the standard R infrastructure. The main in-laboratory developed high-level R functions are available at GitHub repository https://github.com/VGrinev/transcriptome-analysis/blob/master/TranscriptsFeatures.

#### 3.2. Alignment of the short RNA-Seq reads against the reference genome

GRCh38/hg38 reference assembly of the human genome was downloaded as twoBit file from the FTP server of the UCSC Genome Browser. It was then converted to a standard FASTA format with twoBitToFa utility (Speir et al., 2016). A hash table for the reference genome was built with the function *buildindex* from the R/Bioconductor library Rsubread v.1.22.3 (Liao et al., 2013). At this step, 16-mers subreads were extracted in every 3 bases from the reference genome and the threshold 24 was used to exclude highly repetitive subreads from the created hash table.

A global alignment of RNA-Seq reads against the reference genome was carried out with the function *subjunc* from the R/Bioconductor library Rsubread v.1.22.3 (Liao et al., 2013) and the created hash table. This function implements a seed-and-vote mapping paradigm for a fast and accurate alignment (Liao et al., 2013). At this step, we used the default settings of the function *subjunc* and collected only uniquely mapped reads with the maximum of 3 mismatched bases in the alignment. The resulting BAM files were sorted with the function *sortBam* and indexed with the function *indexBam* both from the R/Bioconductor library Rsamtools v.1.24.0 (Morgan, M. et al., 2016).

#### 3.3. Development a list of non-overlapping genomic bins (fragments)

First of all, annotations of the human genes were downloaded from the Ensembl database (see section 2.3). These annotations were converted into an object of a class TranscriptDb (Lawrence et al., 2013) and saved as a local SQLite database. All the subsequent manipulations with genomic intervals were performed using the genomic ranges infrastructure (Lawrence et al., 2013).

Next, the genomic coordinates of the annotated retained introns were calculated as follows. For each gene, the genomic coordinates of exons and introns were extracted from a TranscriptDb object and intersected using the function *findOverlaps* from the R/Bioconductor library GenomicRanges v.1.32.3 (Aboyoun et al., 2018). The intronic intervals that completely fell into the coordinates of exon(-s) of the same gene were selected. These intervals were subsequently intersected with the remaining exons of the gene and, if necessary, disjoined and dropped out due to overlapping. Additionally, the intervals shorter than 100 nucleotides or duplicated intervals were removed from the final list. We called these intervals annotated retained introns, since they are already present in the Ensembl annotation database. Moreover, we further sub-divided these introns into four sub-groups: i) annotated retained introns that completely fall into the alternative first exon (AnnoRI_FIRST), ii) annotated retained introns that are flanked by the internal exons (AnnoRI_INTERNAL), iii) annotated retained introns that completely fall into the alternative last exon (AnnoRI_LAST), and iv) introns that can be inferred as retained because of the overlap with one-exon transcripts (AnnoRI_OneExonTranscript).

Third, the function *intronicParts* of the R/Bioconductor library GenomicFeatures v.1.32.0 (Carlson et al., 2018) was used to extract non-overlapping intronic bins from a TranscriptDb object. Extracted intronic bins were disjoined and dropped out against exons of one- and multi-exons genes (including genes of rRNAs, tRNAs, miRNAs, miscRNAs, ribozymes, vaultRNAs, sRNAs, snRNAs, scaRNAs, scRNAs and snoRNAs). Additionally, any intronic bins shorter than 100 nucleotides were removed. We called these intronic bins canonical introns. The final list of such introns was extended with the annotated retained introns, sorted, indexed, assigned with genes information and converted into an object of a class GRanges.

Fourth, the function *exonicParts* of the R/Bioconductor library GenomicFeatures v.1.32.0 (Carlson et al., 2018) was used to extract non-overlapping exonic bins from a TranscriptDb object. Extracted genomic bins were disjoined and dropped out against genomic coordinates of the annotated retained introns, and genomic bins shorter than 10 nucleotides were removed. The final list of the genomic bins was sorted, indexed, annotated with genes information and converted into an object of a class GRanges.

Finally, all the above mentioned GRanges objects were joined into GRangesList and used in the downstream analysis.

#### 3.4. Development a list of exon clusters

For each gene, the genomic coordinates of exons were retrieved from Ensembl annotations. These coordinates were intersected and joined into overlapping groups called exon clusters. The exon clusters shorter than 10 nucleotides were removed, and the final list of genomic intervals was sorted, assigned with genes information and converted into an object of a class GRanges.

#### 3.5. Read summarisation

We used the function *featureCounts* from the R/Bioconductor library Rsubread v.1.22.3 (Liao et al., 2013, 2014) to assign the mapped RNA-Seq reads to the genomic features in case of exonic and/or intronic bins or to the meta-features (genes) in case of exon clusters. Each read pair was counted in unstranded mode with the minimum 1 base overlapping an exonic bin or exon cluster and minimum 5 bases overlapping an intronic bin. Herewith only one end of the read pair was required to be successfully aligned before the read pair was assigned to a feature or meta-feature.

#### 3.6. *In silico* identification of retained introns

First, the primary count matrix of intronic bins (see sections 3.3 and 3.5) was loaded into the R workspace and an effective length for each intron was calculated using the wgEncodeCrgMapabilityAlign100mer mapability table (Derrien et al., 2012) from the UCSC Genome Browser (Speir et al., 2016). During this step, positions of the non-unique 100-mer alignments and 5 nucleotides from each end of the intron (see section 3.5) were summed and then subtracted from the original intron's length to produce mapability-adjusted intron length.
Second, the primary count matrix of the intronic bins was filtered against the non-expressed genes, intron effective length less than 100 nucleotides and one-bin genes. Third, a variance stabilizing transformation based on the square root of the intron effective length adjusted by the RNA-Seq read length was used to weight individual introns (Boutz et al., 2015; Braun et al., 2017). The sum of intronic reads per gene in each RNA-Seq sample was then partitioned and allocated to each intron proportional to its weight. This led to an *in silico* null model sample, one corresponding to each of the original RNA-Seq samples.
Fourth, differential analysis was carried out to determine introns enriched in the observed reads compared to the *in silico* expected reads (if all the introns within a gene are present at equal levels). We used standard DESeq2 (Love et al., 2014) and edgeR/limma (Ritchie et al., 2015; Robinson et al., 2010) pipelines at this step. Herewith, we discretized the null distributions for the first approach, since DESeq2 uses negative binomial generalized linear modelling, and we loaded the null distributions as it they are for the edgeR/limma pipeline.
Finally, the results of the differential analysis were parsed and filtered. We selected only introns that passed a false discovery rate (FDR) adjusted p-value threshold of 0.01, fold change threshold of 2 and a required minimum of 20 reads per 100 nucleotides of the effective length of an intron averaged over all the original RNA-Seq samples. These introns were called *in silico* detected retained introns, or simply retained introns. The primary count matrix of intronic bins was then reduced to a list of retained introns and it was added to the primary count matrix of the exonic bins.

#### 3.7. Inferring of the differentially used exons (diffUEs)

##### 3.7.1. Identification of diffUEs with DEXSeq

First, the primary count matrix of the exonic bins was loaded into the R workspace and filtered against non-expressed genes, one-bin genes and too low sequencing depth. The filtered count matrix was subsequently used to create a flattened GTF file and it was wrapped (together with a flattened GTF file, sample annotations and experimental design) into an object of a class DEXSeqDataSet (Anders et al., 2012).
Second, the size of each RNA-Seq library was normalized using the "median ratio method" (Anders and Huber, 2010) and dispersion estimates were obtained using the function *estimateDispersions* from the R/Bioconductor library DESeq2 v.1.20.0 (Love et al., 2014; Love et al., 2018). Third, the diffUEs were determined using the functions *testForDEU* and *estimateExonFoldChanges* from the R/Bioconductor library DESeq2 v.1.20.0 (Love et al., 2018) in the default mode. At last, the final results were summarized using the function *DEXSeqResults* from the R/Bioconductor library DESeq2 v.1.20.0 (Love et al., 2018).

##### 3.7.2. Identification of diffUEs with function *diffSplice*

First, the primary count matrix of exonic bins was subjected to filtering against the non-expressed genes, one-bin genes and too low sequencing depth and it was wrapped (together

with the sample information) into a DGEList object (Anders et al., 2013). Second, to calculate effective sizes of RNA-Seq libraries, the scaling factors were estimated using the “trimmed mean of M-values” method (Robinson et al., 2010). Third, by applying the calculated scaling factors, the count data were converted into counts per million, or CPM, and logarithmically transformed, the mean-variance relationship was estimated, and the appropriate observational-level weights were calculated using the voom algorithm (Law et al., 2014).
Fourth, the multiple simple linear models were fitted to the normalized count matrix by least squares method using the function *lmFit* from the R/Bioconductor library limma v.3.36.1 (Ritchie et al., 2015; Smyth et al., 2018). Fifth, contrast coefficients (logarithms for base two of fold changes, or $\log_2$ FC, between the treatment conditions) were calculated and loaded into the function *diffSplice* (Ritchie et al., 2015; Smyth et al., 2018) . This function calculates the difference between the $\log_2$ FC for a given exon versus the average $\log_2$ FC for all the other exons for the gene of interest. In other words, this function tests for differential usage of exons for each gene and for each treatment condition. Finally, from moderated t-statistics, p-values were adjusted for multiple testing with the method by Benjamini Y. and Hochberg Y., which controls the expected FDR below the specified value (Benjamini and Hochberg, 1995).

##### 3.7.3. Identification of diffUEs using the functionality of the JunctionSeq library

First, the overall quality of the BAM files was assessed with Picard v.2.9.0 (http://broadinstitute.github.io/picard/) and low-quality reads were removed using in-laboratory developed R code. Second, the flattened GFF file was created using toolset QoRTs (Hartley and Mullikin, 2015). This file was based on the Ensemble annotations of the human genome and included all the exons, annotated and novel exon-exon junctions (EEJs). Third, reads counts were generated by QoRTs. At this step, we counted all the reads mapped to exons, annotated or novel EEJs with minimum mapping quality of 30.
Fourth, the diffUEs were identified by the sequential application of two functions *runJunctionSeqAnalyses* and *writeCompleteResults* in the default mode to reads counts. These functions are part of the R/Bioconductor library JunctionSeq v.1.10 (Hartley and Mullikin, 2016) and they use DEXSeq statistical infrastructure (Anders et al., 2012) to detect diffUEs. Finally, output results were parsed and adjusted to the formats of DEXSeq *and diffSplice* outputs by in-laboratory developed R code.

#### 3.8. Functional classification of exons

Genomic coordinates of the reference exons were extracted from the Ensembl models of the human genes. These exons were grouped into five functional classes: 5’UTR exons, CDS exons, 3’UTR exons, exons of non-coding RNAs (NC) and multi-type exons (MTE) (exons that can be non-coding, 5’UTR, 5’UTR/CDS, CDS, CDS/3’UTR and/or 3’UTR exon depending on transcript). Next, each exonic bin from section 3.3 was intersected with reference exons and was assigned to a functional class.

#### 3.9. Identification of EEJs

All possible variants of EEJs were identified according to Liao et al. (Liao et al., 2013). The resulting BED files were parsed and converted into the primary count matrix of EEJs with an in-laboratory developed R code. This matrix included a full list of identified EEJs with the respective genomic coordinates and a number of reads supporting each exon-exon junction in every sample.

#### 3.10. Inferring of the differentially used exon-exon junctions (diffEEJs)

##### 3.10.1. Identification of diffEEJs with function *diffSplice*

The primary count matrix of EEJs was subjected to filtering against too low sequencing depth and it was wrapped (together with the sample information) into a DGEList object (Anders et al., 2013). All the subsequent steps of the analysis were carried out in accordance with subsection 3.7.2, but at the level of EEJs.

##### 3.10.2. Identification of diffEEJs using functionality of JunctionSeq library

Inferring of diffEEJs using the JunctionSeq library was performed as described in subsection 3.7.3, but at the level of EEJs.

#### 3.11. Classification of EEJs according to the modes of alternative splicing

Our classifier of EEJs is based on the idea of hypothetical "non-alternative" precursor of RNA, or hnapRNA. hnapRNA is an RNA molecule that would have turned out if the gene had only one transcription start site (TSS), if there were no alternative splice sites, if there was no alternative splicing and if there was only one transcription termination site. In other words, hnapRNA is a generalization of all the RNA isoforms produced by the gene.

For each gene, the structure of the hnapRNA was calculated using Ensembl models of human genes. We clustered exons of the gene of interest into overlapping groups with the exception of retained introns, alternative 5’ and/or 3’ terminal exons. The outer boundaries of the resulting exon clusters were recorded as genomic coordinates of exons of the hnapRNA. The list of these coordinates was extended with coordinates of retained introns, alternative 5’ and/or 3’ terminal exons and it was converted into an object of a class GRanges.

Next, the genomic coordinates of EEJs were intersected with the coordinates of the features of the hnapRNA, and the mode of each EEJ was determined. According to our approach, all the EEJs were classified into eight modes of alternative splicing:

i) canonical event, if the coordinates of the empirical event exactly match the model event,

ii) alternative 5' splice site, if only the 3’ splice site of the empirical event exactly matches the respective model site,

iii) alternative 3' splice site, if only the 5’ splice site of the empirical event exactly matches respective model site,

iv) alternative both splice sites (intron isoform), if both splice sites of the empirical event do not match splice sites of respective model event,

v) skipped cassette exon(-s), if the empirical event includes skipping one or more exons of the model,

vi) alternative first exon, if the 5’ splice site of the empirical event exactly matches the 3’ end of alternative first exon in model,

vii) alternative last exon, if the 3’ splice site of the empirical event exactly matches the 5’ end of alternative last exon in model,

viii) complex splicing event, if the empirical event includes two or more of the above-mentioned alternative splicing events.

#### 3.12. Reference-based transcriptome assembly

First, for each sample of RNA, we used Cufflinks (Trapnell et al., 2010) and the respective *subjunc*-generated BAM file to assemble the alignments into a parsimonious set of transcripts. Herewith, Cufflinks was supplied with i) Ensembl annotation of the human genome to guide RABT assembly, ii) a GTF file containing annotated human rRNA and mitochondrial genes to mask these genomic features during estimation of transcripts abundance, iii) complete sequence of the human genome in multiFASTA format to bias correction during the estimation of transcripts abundance, and iv) a minimal isoform fraction threshold assigned to 0.05.

Second, individual Cufflinks assembled transcriptomes were merged into one consolidated set of transcripts with Cuffmerge (Trapnell et al., 2012). This set of transcripts was filtered against i) unstranded transcripts, ii) too short transcripts (<300 nucleotides), iii) transcripts with too short exon(-s) (<25 nucleotides), iv) transcripts with too short intron(-s) (<50 nucleotides), and vi) transcripts with low abundance (fragments per kilobase of transcript per million mapped reads, or FPKM, below 1). Filtration was controlled by in-laboratory developed R code.

Third, the consolidated and filtered set of transcripts was submitted to Cuffdiff (Trapnell et al., 2013) for the simultaneous calculation of the transcript abundance and differential expression. Cuffdiff was provided with a GTF file containing annotated human rRNA and mitochondrial genes and a multiFASTA file with complete sequence of the human genome, and it was run in default mode except for the minimal isoform fraction threshold that was assigned to 0.05. Finally, for the fast retrieving of the data and easy subsequent manipulations, the main outcomes of Cuffdiff were parsed, converted into an object of a class TranscriptDb (Lawrence et al., 2013) and saved as a local SQLite database.

### 3.13. Analysis of differential gene expression

#### 3.13.1. Identification of differentially expressed genes with Cuffdiff

We used Cuffdiff differential expression tests data (see section 3.12 above) to identify differential expression at transcript or gene levels between short interfering RNA treatment conditions. Herewith, only transcripts or genes with at least 2-fold changes in expression and q-value below 0.1 were annotated as differentially expressed.

#### 3.13.2. Identification of differentially expressed genes with DESeq2

First, the mapped RNA-Seq reads were assigned to the genomic meta-features (genes) as described in section 3.5. Second, the resulting count matrix was subjected to filtering against too low sequencing depth and it was wrapped (together with the sample information) into a DESeqDataSet object (Love et al., 2018). Third, differentially expressed genes were identified using the functions *DESeq* and *results* from the R/Bioconductor library DESeq2 v.1.16.1 (Love et al., 2018). These functions were run in default mode and according to the standard DESeq2 pipeline. Finally, results were parsed and genes with at least 2-fold changes in expression and q-value below 0.1 were annotated as differentially expressed.

#### 3.13.3. Identification of differentially expressed genes with edgeR/limma

First, the mapped RNA-Seq reads were assigned to the genomic meta-features as described in section 3.5. Second, the resulting count matrix was subjected to filtering against too low sequencing depth and it was wrapped (together with the sample information) into a DGEList object (Anders et al., 2013). Third, to calculate an effective size of each RNA-Seq library, the scaling factors were estimated using the “trimmed mean of M-values” method (Robinson et al., 2010). Fourth, by applying the calculated scaling factors, the count data were converted into CPM and logarithmically transformed, the mean-variance relationship was estimated, and the appropriate observational-level weights were calculated using the voom algorithm (Law et al., 2014).

Fifth, the multiple simple linear models were fitted to the normalized count matrix by least squares method using the function *lmFit* from the R/Bioconductor library limma v.3.36.1 (Ritchie et al., 2015; Smyth et al., 2018). Sixth, $\log_2$ FC coefficients and empirical Bayes statistics were calculated using respective functions from R/Bioconductor libraries edgeR v.3.22.3 (Chen et al., 2018; Robinson et al., 2010) and limma v.3.34.9 (Ritchie et al., 2015;

Smyth et al., 2018). Finally, results were parsed and genes with at least 2-fold changes in expression and the q-value below 0.1 were annotated as differentially expressed.

##### 3.13.4. Identification of differentially expressed and differentially used TSSs

Differentially expressed TSSs were identified according to the Cuffdiff algorithm (Trapnell et al., 2013). Differential usage of TSSs was analysed according to idea implemented in *diffSplice* function of the R/Bioconductor library limma v.3.34.9 (Ritchie et al., 2015; Smyth et al., 2018) for each gene containing alternative TSSs.

#### 3.14. Linear approximation of datasets

##### 3.14.1. Linear approximation using a principal component analysis (PCA)

A standard PCA on the given data matrix was performed using the basic R function *prcomp*. This function was provided with centred and scaled data. Alternatively, multigroup PCA and/or kernel PCA were carried out using the R library “multigroup” v.0.4.4 (Eslami et al., 2015) and “kernlab” v.0.9-26 (Karatzoglou et al., 2004), respectively. The results of the linear approximations were used to calculate the “explained” variability and visualisation of the multidimensional data in the space (usually) of the first two principal components.

##### 3.14.2. Linear approximation using a t-distributed stochastic neighbor embedding (t-SNE)

A t-SNE was used as an alternative to the PCA where a deeper linear approximation of the data matrix was necessary. We used an R wrapper Rtsne (Krijthe and van der Maaten, 2017) for the Van der Maaten’s C++ implementation of the Barnes-Hut algorithm of t-SNE (Maaten, 2014). This wrapper was run in the default mode except for perplexity and theta values, which were sequentially adjusted.

##### 3.14.3. Linear approximation using an independent component analysis (ICA)

An ICA was run on the combined microarray dataset for unsupervised separation of cell types and extraction of cell specific genes. To perform analysis, an R wrapper fastICA (Marchini et al., 2017) for the Aapo Hyvärinen’s implementation of the FastICA algorithm (Hyvärinen and Oja, 2000) was used in the default mode. The set of the most significant genes per independent component was detected at FDR < 0.05 as described previously (Nazarov et al., 2018).

#### 3.15. Identification of direct targets of RUNX1/RUNX1T1 fusion protein

We reanalysed our previous ChIP-Seq data to identify RUNX1/RUNX1T1 binding peaks in the genome of Kasumi-1 cells (Ptasinska et al., 2012). We used GSM722718, GSM722706 and GSM722707 FASTQ files for input, siMM- and siRR-treated Kasumi-1 cells. Reads were aligned against GRCh38/hg38 reference assembly of the human genome as described in section 3.2. At this step, the function *align* was used instead of the function *subjunc* from the R/Bioconductor library Rsubread v.1.22.3 (Liao et al., 2013).

ChIP-Seq peaks were called in a frame of a fully Bayesian hidden Markov model with 10000 Monte Carlo runs for each Markov chain (Cairns et al., 2011). Next, the peaks with more than 99.9% posterior probabilities were selected and depth/coverage filtered. The genomic coordinates of the final peaks were intersected with coordinates of the human genes and direct targets were identified. A gene was annotated as a direct target of the fusion protein if it has RUNX1/RUNX1T1 peak(-s) within its transcription unit or in the immediate vicinity, < 3000 bp upstream of the transcription start site.

#### 3.16. Development a list of features associated with EEJs

Every exon-exon junction was annotated with sequence, sequence-related, functional, and structural features that were extracted from four types of genomic/RNA elements: 100-bp fragment of the upstream exon (USE), 300-bp fragment from the 5’ end of the intron (USIF), 300-bp fragment from the 3’ end of the intron (DSIF), and 100-bp fragment of the downstream exon (DSE). Additionally, each exon-exon junction was described with a set of nearest epigenetic marks. In total, the complete list of features included 1680 items.

#### 3.16.1. Sequence features

##### *Splice sites scoring*

Genomic coordinates of the 5’ and 3’ splice sites were retrieved from the matrix of the experimentally identified EEJs (see section 3.9) and sequences of these sites were extracted from the GRCh38/hg38 reference assembly of the human genome with the R/Bioconductor library BSgenome.Hsapiens.UCSC.hg38 (Team, 2015). Position weight matrices for the 5’ splice sites (9-nucleotide sequence – 3 nucleotides in the exon and 6 nucleotides in the intron) and 3’ splice sites (23-nucleotide sequence – 3 nucleotides in the exon and 20 nucleotides in the intron) were downloaded from the MIT MaxEnt Splice Site Scoring Server (Yeo et al., 2004). The strength of splice sites was determined by Perl implementation of the MaxEntScan algorithms in accordance with the three scoring models: maximum entropy model, first-order Markov model, and weight matrix model. The overall score of the splice sites for a given exon-exon junction was calculated as the sum of individual 5’ splice site and 3’ splice site scores averaged over the three models.

##### *Experimentally verified exonic splicing motifs*

We collected sequences of experimentally verified binding sites that were recognized by DAZAP (Goina et al., 2008; Haque et al., 2010; Skoko et al., 2008), ELAVL1 (Doller et al., 2010; Katsanou et al., 2005; Ma et al., 1996; Woo et al., 2009), FMR1 (Didiot et al., 2008; Schaeffer et al., 2001), HNRNPA1 (Burd and Dreyfuss, 1994), HNRNPA2B1 (Goina et al., 2008; Haque et al., 2010; Skoko et al., 2008), HNRNPC (Hamilton et al., 1993; Kim et al., 2003; Temsamani and Pederson, 1996), HNRNPD (Chang et al., 2010; Chen et al., 2001; Fialcowitz et al., 2005; Palanisamy et al., 2008; Pullmann et al., 2006; Sommer et al., 2005), HNRNPL (Heiner et al., 2010; Motta-Mena et al., 2010; Rothrock et al., 2005), HNRNPLL (Preußner et al., 2012; Temsamani and Pederson, 1996; Topp et al., 2008), HNRNPU (Haque et al., 2010), SRSF1 (Cartegni et al., 2003, 2006; Smith et al., 2006), SRSF2 (Cartegni et al., 2003, 2006; Smith et al., 2006), SRSF3 (Galiana-Arnoux et al., 2003; Gonçalves et al., 2009; Jang et al., 2014), SRSF5 (Cartegni et al., 2003, 2006; Smith et al., 2006), SRSF6 (Cartegni et al., 2003, 2006; Smith et al., 2006), and TARDBP (Ayala et al., 2006; Buratti et al., 2010; Dujardin et al., 2010; Mercado et al., 2005; Volkening et al., 2009) splicing-related proteins. For each protein and the respective set of sequences, we performed a motif search by the discriminative motif discovery algorithm motifRG (Yao et al., 2014) against the background set of randomly extracted human intronic and exonic sequences. The primary motif was refined by the function *refinePWMMotif* from the R/Bioconductor library motifRG v.1.18.0 (Yao et al., 2014) with the default settings and was converted into the $\log_2$ position weight matrix with the correction against the background nucleotides frequency. Occurrence of a motif in the USE and DSE was determined by the function *countPWM* from the R/Bioconductor library Biostrings v.2.42.0 (Pages, H et al., 2014) and was normalized relative to the length of the analysed sequences. We used the $99^{th}$ quantile of the motif weight distribution as a threshold in the identification of the true motif occurrence.

Additionally, we collected oligomeric sequences that were bound by splicing proteins ELAVL4 (Chung et al., 1996; Joseph et al., 1998; Zhu et al., 2006), HNRNPA3 (Chen et al., 2008), HNRNPC2 (Haque et al., 2010; Pullmann et al., 2006), HNRNPC1 (Pullmann et al., 2006), HNRNPDL (Zubović et al., 2012), HNRNPF (Dominguez and Allain, 2006; Mauger et al., 2008; Millevoi et al., 2009), HNRNPH1 (Fisette et al., 2010; Millevoi et al., 2009; Ohe et al., 2010; Russo et al., 2010), HNRNPH2 (Fisette et al., 2010; Masuda et al., 2008; Millevoi et al., 2009; Ohe et al., 2010), HNRNPH3 (Masuda et al., 2008; Millevoi et al., 2009; Ohe et al., 2010), HNRNPK (Lee et al., 2007; Melton et al., 2007; Motta-Mena et al., 2010), HNRNPM

(Cho et al., 2014), KHDRBS1 (Itoh et al., 2002; Pedrotti et al., 2010; Sellier et al., 2010), KHSRP (García-Mayoral et al., 2007; Linker et al., 2005; Trabucchi et al., 2009; Winzen et al., 2007), MBNL1 (Goers et al., 2010; Ho et al., 2004; Sellier et al., 2010; Sen et al., 2010), PCBP1 (Holcik and Liebhaber, 1997; Lee et al., 2007; Rothrock et al., 2005; Thomson et al., 2005), PCBP2 (Eiring et al., 2010; Motta-Mena et al., 2010; Rothrock et al., 2005; Thomson et al., 2005), QKI (Zong et al., 2014), RBM25 (Zhou et al., 2008), SF3B1 (Massiello et al., 2006), SFPQ (Buxadé et al., 2008; Hall-Pogar et al., 2007; Melton et al., 2007; Motta-Mena et al., 2010), SRP54 (Wu et al., 2006), SRSF4 (Buratti et al., 2004), SRSF7 (Galiana-Arnoux et al., 2003; Schaal and Maniatis, 1999; Venables et al., 2005), SRSF9 (Cloutier et al., 2008; Simard and Chabot, 2002), SYNCRIP (Chen et al., 2008; Duning et al., 2008), TIA1/TIAL1 (Aznarez et al., 2008; Buxadé et al., 2008; Izquierdo and Valcárcel, 2007; McAlinden et al., 2007; Reyes and Izquierdo, 2007; Wang et al., 2014; Zhu et al., 2006) and YBX1 (Fraser et al., 2008; Shen et al., 2006; Skoko et al., 2008; Wei et al., 2012). We were not able to calculate position weight matrices of the motifs for these proteins due to a limited number of sequences of the experimentally verified binding sites. For this reason, we used an alternative approach in the assessment of the strength of binding sites (BSS) for the mentioned above splicing proteins, as proposed by Murray et al. (Murray et al., 2008):

$$BSS = \frac{\sum_{i=0}^{i=L-k+1} ln(4^k f_{n_i})}{L-k+1},$$

where *L* is the length of the sequence of interest, *k* is the length of an oligomer (see below) found in the sequence of interest, $f_n$ represents the frequency (within the set of sequences of the experimentally verified binding sites for a given splicing protein) of the oligomer found at the position *i* in the sequence of interest, and $ln(4^k f_{n_i})$ is a log-odds representation of the degree to which the particular oligomer was enriched within the set of sequences of the experimentally verified binding sites for a given splicing protein. As proposed, we counted only the frequency of all the possible pentamers in the sequence of interest and used the frequency of pentamers from the set of sequences of the experimentally verified binding sites for a given splicing protein as the reference (Murray et al., 2008).

***Bioinformatically predicted exonic splicing motifs***

We selected three different approaches for *de novo* motifs discovery and identified 25 new motifs that were statistically associated with multi-spliced exons from the Ensembl database. First of all, we used the algorithm GADEM (Li, 2009) from R/Bioconductor library rGADEM v.2.22.0 (Droit et al., 2014). This approach was realized on a sub-set of multi-spliced exons with default settings of the software. The second approach was based on the heuristic algorithm bcrank from R/Bioconductor library BCRANK v.1.36.0 (Ameur et al., 2009; Ameur, 2010). In this case, short sequences that were overrepresented in ranked exons (in descending order of their splicing degrees) were identified and the top motifs were selected for subsequent analysis. Finally, the algorithm motifRG from the R/Bioconductor library motifRG v.1.18.0 (Yao et al., 2014) was used with default settings. This algorithm searches for motifs that discriminate the given foreground and background sequences. We used a sub-set of multi-spliced exons as foreground sequences and other exons from our dataset as background sequences.

All newly identified motifs were converted into $log_2$ position weight matrices with the correction against the background nucleotide frequency. Position weight matrices of Sironi's motifs 1-3 (Sironi et al., 2004) were added to our collection of bioinformatically predicted exonic splicing motifs. Occurrence of these motifs in the sequence of interest was determined as described above. In addition, oligomers with the bioinformatically predicted exonic splicing activity were counted in the USE and DSE by the function *vcountPDict* from the

R/Bioconductor library Biostrings v.2.42.0 (Pages et al., 2014) and their frequency was normalized relative to the length of the analysed sequences. The list of such oligomers included ESRE hexamers (Goren et al., 2006), ESRS hexamers (Goren et al., 2006), ESS decamers (Wang et al., 2004), PESE octamers (Zhang and Chasin, 2004; Zhang et al., 2005), PESS octamers (Zhang and Chasin, 2004; Zhang et al., 2005), QUEPASA ESEseqs hexamers (Ke et al., 2011), QUEPASA ESSseqs hexamers (Ke et al., 2011), and RESCUE ESE hexamers (Fairbrother et al., 2002).

***Experimentally verified intronic splicing motifs***

We could reconstruct the position weight matrices of the motifs for seven splicing proteins that bind intronic sequences: CELF1 (Dujardin et al., 2010; Le Tonquèze et al., 2010; Marquis et al., 2006; Vlasova et al., 2008), HNRNPA1 (Burd and Dreyfuss, 1994), HNRNPA2B1 (Goina et al., 2008; Haque et al., 2010; Skoko et al., 2008), HNRNPL (Heiner et al., 2010; Motta-Mena et al., 2010; Rothrock et al., 2005), SRSF3 (Galiana-Arnoux et al., 2003; Gonçalves et al., 2009; Jang et al., 2014; Lou et al., 1998), TARDBP (Ayala et al., 2006; Buratti et al., 2010; Dujardin et al., 2010; Mercado et al., 2005; Volkening et al., 2009) and TRA2B (Cho et al., 2014; Venables et al., 2005; Wu et al., 2006). We also collected the experimentally verified intronic sequences that are bound by splicing proteins CELF3 (Dujardin et al., 2010), ELAVL2 (Zhu et al., 2006), ELAVL4 (Chung et al., 1996; Joseph et al., 1998; Zhu et al., 2006), HNRNPA3 (Chen et al., 2008), (Zubović et al., 2012), HNRNPF (Dominguez and Allain, 2006; Mauger et al., 2008; Millevoi et al., 2009), HNRNPH1 (Fisette et al., 2010; Millevoi et al., 2009; Ohe et al., 2010; Russo et al., 2010), HNRNPH2 (Fisette et al., 2010; Masuda et al., 2008; Millevoi et al., 2009; Ohe et al., 2010), HNRNPH3 (Masuda et al., 2008; Millevoi et al., 2009; Ohe et al., 2010), MBNL1 (Goers et al., 2010; Ho et al., 2004; Sellier et al., 2010; Sen et al., 2010), PTBP1 (David et al., 2010; Han et al., 2014; Motta-Mena et al., 2010; Zubović et al., 2012), PTBP2 (Zubović et al., 2012), QKI (Zong et al., 2014), RBFOX1 (Ponthier et al., 2006; Zhou et al., 2008, 2007; Zubović et al., 2012), RBFOX2 (Ponthier et al., 2006; Zhou et al., 2008, 2007), RBFOX3 (Kim et al., 2003; Minovitsky et al., 2005), RBM4 (Kar et al., 2006; Lin and Tarn, 2005), SF1 (Královicová et al., 2004; Zong et al., 2014), SRSF7 (Galiana-Arnoux et al., 2003; Schaal and Maniatis, 1999; Venables et al., 2005), SRSF9 (Cloutier et al., 2008; Simard and Chabot, 2002), SYNCRIP (Chen et al., 2008; Duning et al., 2008), TIA1/TIAL1 (Aznarez et al., 2008; Buxadé et al., 2008; Izquierdo and Valcárcel, 2007; McAlinden et al., 2007; Reyes and Izquierdo, 2007; Wang et al., 2014; Zhu et al., 2006), TRA2A (Tacke et al., 1998) and YBX1 (Fraser et al., 2008; Shen et al., 2006; Skoko et al., 2008; Wei et al., 2012), but for which we could not calculate the position weight matrices because of a limited number of sequences. We used the described above approaches for the determination of occurrence of all these motifs in the USIF and DSIF. It should be noted that some splicing proteins did not exhibit any intron/exon preference and are bound to both intronic and exonic motifs.

***Bioinformatically predicted intronic splicing motifs***

Oligomers with the bioinformatically predicted splicing activity were counted in the USIF and DSIF as described above. The list of such oligomers included Castle's oligomers (Castle et al., 2008), Culler's ISSs oligomers (Culler et al., 2010), Das' upstream intronic hexamers (Das et al., 2007) , Das' downstream intronic hexamers (Das et al., 2007), Wang's ISEs hexamers (Wang et al., 2012), Wang's ISSs decamers (Wang et al., 2013), Yeo's downstream ISREs (Yeo et al., 2007) , and Yeo's upstream ISREs (Yeo et al., 2007).

***Polypyrimidine tract scoring***

For each splicing event, the sequence of the polypyrimidine tract (-30 to -3 positions relative to the acceptor splice site) was extracted from the corresponding DSIF. The strength of U2AF2 binding sites in this sequence was calculated according to Murray et al. (Murray et al., 2008):

$$U2AF2_{strength} = \frac{\sum_{i=0}^{i=L-k+1} ln(4^k f_{n_i})}{L-k+1},$$

where *L* is the length of the extracted polypyrimidine tract sequence, *k* is the length of an oligomer found in the polypyrimidine tract sequence, $f_n$ represents the frequency (within the U2AF2 selected SELEX sequences) of the oligomer found at the position *i* in the polypyrimidine tract sequence and $ln(4^k f_{n_i})$ is the log-odds representation of the degree to which a particular oligomer was enriched within the U2AF2 selected SELEX sequences (Banerjee et al., 2004). This equation is identical to the equation for BSS, however, the values of independent variables in this equation allow us to work only with binding sites for the protein U2AF2. We counted the frequency of all the possible pentamers in the polypyrimidine tract and used the frequency of pentamers from the U2AF2 selected SELEX sequences as the reference (Murray et al., 2008).

Moreover, we collected G- and C-rich 4- to 7-nucleotide sequences overrepresented in intronic regions upstream of the weak polypyrimidine tracts (Murray et al., 2008). We determined the frequency of these motifs in DSIF (-80 to -30 positions relative to the acceptor splice site) using the function *vcountPDict* from the R/Bioconductor library Biostrings v.2.42.0 (Pages et al., 2014) and normalized this parameter relative to the length of the analyzed intronic fragment.

***Branchpoint sites scoring***

First of all, we obtained genomic coordinates of 59359 high-confidence human branchpoint sites from the work by Mercer et al. (Mercer et al., 2015). Next, we retrieved the 20-nucleotide sequences surrounding the branchpoints from the GRCh38/hg38 reference assembly of the Homo sapiens genome using the R/Bioconductor library BSgenome.Hsapiens.UCSC.hg38 (Team, 2015) and used these sequences as a foreground set in the motif discovery analysis. Additionally, we developed a background set of sequences with 10-fold quantitative excess relative to the foreground set. This background set included randomly extracted 100-nucleotide sequences from the upstream regions of human introns.

Next, we used the discriminative motif discovery algorithm motifRG with default settings (Yao et al., 2014) and identified high confidence motif associated with human branchpoint sites. This motif was converted into $log_2$ position weight matrix with the correction against the background nucleotide frequency and used for the scanning of the sequence of interest. For each splicing event, we extracted a 100-nucleotide sequence (-100 to -1 positions relative to the acceptor splice site) from the corresponding DSIF. For each extracted sequence, we calculated the maximal and mean motif affinity to the sequence of interest and a number of hits over the score threshold using the function *motifScores* from the R/Bioconductor library PWMEnrich v.4.10.0 (Stojnic and Diez, 2015). As before, we used the $99^{th}$ quantile of the motif weight distribution as a threshold in the identification of the true motif occurrence.

***"Short" motifs***

Frequency of *x*-mer (at $x \in [1,4]$) oligonucleotides in the sequence of interest was determined using the function *oligonucleotideFrequency* from the R/Bioconductor library Biostrings v.2.42.0 (Pages et al., 2014) and normalized relative to the length of the analyzed sequence.

**3.16.2. Sequence-related features**

***Linear density of the minimal free energy of folding***

The sequences of USE, USIF, DSIF, or DSE genomic/RNA elements were extracted from the GRCh38/hg38 reference assembly of the *Homo sapiens* genome using the R/Bioconductor library BSgenome.Hsapiens.UCSC.hg38 (Team, 2015). Free energy of folding, or minimal free energy (MFE), of these sequences was calculated using the RNAfold tool from ViennaRNA Package v.2.1.7 (Lorenz et al., 2011). MFE was normalized relative to the length of the analyzed sequence and expressed as a linear density of the MFE (Pervouchine et al., 2003; Posrednik et al., 2011).

***Conservation scores***

BW files with pre-computed conservation scores of the human GRCh38/hg38 reference genome were downloaded via the FTP server of the UCSC Genome Browser (Harrow et al., 2014). We used conservation scores that were calculated using the algorithms phyloP and phastCons after multiz-based multiple alignments of 99 vertebrate genomes to the human genome (Pollard et al., 2010; Siepel et al., 2005). Using the downloaded BW files, we calculated the minimum, maximum, standard deviation, and mean scores for each sequence of interest.

#### 3.16.3. Functional features

Each splice site of EEJs was intersected with genomic coordinates of the splice sites of functionally grouped exons (see section 3.8). Next, exactly matched splice sites of EEJs were assigned to a functional class of respective exon.

#### 3.16.4. Structural features

***Splicing distances***

Splicing distances (length of introns) were directly retrieved from the matrix of the experimentally identified EEJs (see section 3.9).

***Size of exon clusters***

This metric describes a distribution of constitutive and alternative splice sites along the body of the gene in the immediate vicinity of the splice site of interest. Exon clusters were calculated as described in section 3.4. The number of exons in a cluster was considered as the size of the cluster. Each splice site of EEJs was assigned the size of the exon cluster to which it and its partner splice site(-s) belonged.

#### 3.16.5. Epigenetics features

In this study, we used data describing the location of five epigenetics marks in the genome of the siRR- or siMM-treated Kasumi-1 cells: CpGs islands, DNase I hypersensitivity sites, modified histone H3K9Ac, RNA polymerase II peaks and RUNX1/RUNX1T1 peaks (see sections 2.1 and 2.3). Distances of the splice sites of EEJs to the nearest epigenetic marks were measured using the function *distanceToNearest* from the R/Bioconductor library GenomicRanges v.1.32.3 (Aboyoun et al., 2018).

### 3.17. Data mining with the random forest meta-classifier

#### 3.17.1. Filtration of the primary data matrix

Our primary data matrix included the class of exon-exon junction (differential or non-differential) as a dependent response variable and all the features described above as an independent predictor of variables. This matrix was filtered against the features that had only one unique value or features that had both characteristics: i) very few unique values relative to the number of samples and ii) the ratio of the frequency of the most common value to the frequency of the second most common value is large. Highly correlated features were removed with a cut-off 0.9 to reduce pair-wise correlations in the matrix. At this step, we used the functionality of the R library caret v.6.0-71 (Kuhn, 2016).

#### 3.17.2. Feature importance

The importance of each feature was determined by calculating the total decrease in the node impurities from splitting on the feature averaged over all classification trees in random forest. The node impurity was measured with the Gini index. At this step, we used the R library randomForest v.4.6-12 in the classification mode (Breiman et al., 2016; Liaw and Wiener, 2002). We also used five independent runs of the random forest meta-classifier, and 1000 classification trees per random forest per run and ranked all features in descending order of importance.

**3.17.3. Feature selection**

We applied a recursive algorithm with five-fold cross-validation to select the minimal required set of important features. We used the function *rfe* from the R library caret v.6.0-71 at this step (Kuhn, 2016).

**3.17.4. Final classification of EEJs**

First of all, the data matrix was reduced to features selected in the previous step. Next, for each run of the random forest meta-classifier, the data matrix was randomly sampled on two sub-matrices: the training set (70% of input matrix) and the test set (30% of input matrix). We used the training set for machine learning, calculation of the proximity matrix and marginal effects of features. The test set was used to assess the classification accuracy. We used the R library randomForest v.4.6-12 in the classification mode at this step (Breiman et al., 2016; Liaw and Wiener, 2002), 1000 trees were grown at each algorithm run and the number of features sampled for splitting up at each node was equal to one third of all features in the input data matrix.

### 3.18. Reconstruction of the gene-regulatory networks

The sequences of promoter regions of the genes of interest were scanned against 2287 position weight matrices of the human transcription factors from the MotifDb database (Shannon, 2018). To find out the true motifs in promoter sequences, lognormal threshold-free approach was used instead of the fixed-threshold algorithm. In this instance, transcription factors with significantly enriched motifs in the promoters of interest (enrichment score > 1 at $p < 0.05$ compared to a genomic background) and with statistically significant differential expression ($p < 0.01$, FDR-adjusted $p < 0.1$) in the siRR- versus siMM-treated Kasumi-1 cells were selected for downstream analysis. Direct interactions of the fusion protein with the transcription factors genes and the target genes of interest were inferred from the ChIP-Seq data. Co-expression of *RUNX1*/*RUNX1T1*, genes coding transcription factors selected in the second step and the target genes of interest (genes encoding splicing factors and mRNA surveillance genes with differential expression) was inferred from the RNA-Seq and microarray data with CoExpress software (Nazarov et al., 2013). From this analysis, only pairs of genes with more than 90% robust correlation in expression were selected. Genes with any discordance in expression and/or correlations between RNA-Seq and microarray data were excluded from the final list. In this instance, FDR of the correlations did not exceed 6.5% for the RNA-Seq data and 2.9% for the microarray data.

### 3.19. Enrichment tests

**3.19.1. Gene enrichment analysis**

We used up-to-date ODO and GAF files from the Gene Ontology Consortium (Ashburner et al., 2000; The Gene Ontology Consortium, 2017) to develop a comprehensive list of the reference functional gene sets. From this list, we selected the gene sets containing ten or more members for downstream analysis. Next, two-sided Fisher's exact test was used to find out the under- and/or over-represented query gene set(-s) among the reference gene sets. Query results were parsed and under- or over-represented gene sets that passed members size $\geq 10$ and FDR adjusted p-value threshold of 0.05 were collected. Finally, Cytoscape plug-in EnrichmentMap (Merico et al., 2010) was used to handle gene-set redundancy and hierarchical visualization of the enrichment results.

**3.19.2. Gene set enrichment analysis**

Gene set enrichment analysis was performed according to standard procedure using Molecular Signatures Database (Subramanian et al., 2005).

#### 3.20. Assessment of the coding potential of RNA transcripts

Coding potential of RNA transcripts was assessed by on-line version of CPC2 software (Kang et al., 2017). This software uses sequence features of transcripts and support vector machine for reliable prediction of coding ability of RNA molecules (Kong et al., 2007).

#### 3.21. Identification of the significant open reading frames (ORFs) and premature termination codons (PTCs) in transcripts

Significant ORFs and PTCs were identified in the Cufflinks assembled transcripts as previously described (Grinev et al., 2015). In brief, all possible ATG-ORFs were identified in the transcript(s) of interest. Next, for each empirical transcript, 100 random sequences with the same length were generated using a multinomial model (Ababneh et al., 2006). This new set of artificial transcripts was used to identify the ORFs. Finally, the 99th percentile of the distribution formed by the lengths of the artificial ORFs was used as a threshold for identification of the true ORF(s) in the empirical transcript. Transcripts with no significant ORFs were classified as non-coding. To identify PTCs, exonic structure and the coordinates of ORF(s) in the transcript of interest were matched. A transcript was annotated as PTC-containing, if the end of its ORF was localized $\geq$50 nucleotides upstream of the last exon-exon junction in the transcript.

#### 3.22. Alignment classification of the *in silico* translated proteins

The *in silico* translated proteins were aligned against the human up-to-date NCBI RefSeq proteins and the newest non-redundant releases of GenBank CDS translations, UniProtKB/SwissProt, Protein Data Bank, Protein Information Resource and Peptide/Protein Sequence Database proteins using NCBI blastp (Gish et al., 1993). The *in silico* translated proteins with no alignment for any canonical proteins were classified as no hits. Next, according to identity value, all “with hits” proteins were divided into lowly identical (with identity below 90%) and highly identical (identity above 90%) to canonical proteins. Finally, all highly identical proteins were further classified into normal, N-extended, N-truncated, C-truncated, C-extended and complex based on the following criteria: i) normal, the *in silico* translated and reference proteins align perfectly; ii) N-extended, the *in silico* translated protein contains novel N-terminal amino acids followed by the full-length canonical protein sequence; iii) N-truncated, the *in silico* translated protein that lacks N-terminal part of the canonical protein; iv) C-truncated, the *in silico* translated protein that lacks C-terminal part of the canonical protein; v) C-extended, the *in silico* translated protein that includes the full-length canonical protein sequence, followed by the novel C-terminal amino acids; vi) complex, the *in silico* translated protein other than canonical protein at both ends.
