## Supplemental Figures for "RUNX1/RUNX1T1 controls alternative splicing in the t(8;21)-positive acute myeloid leukemia cells"

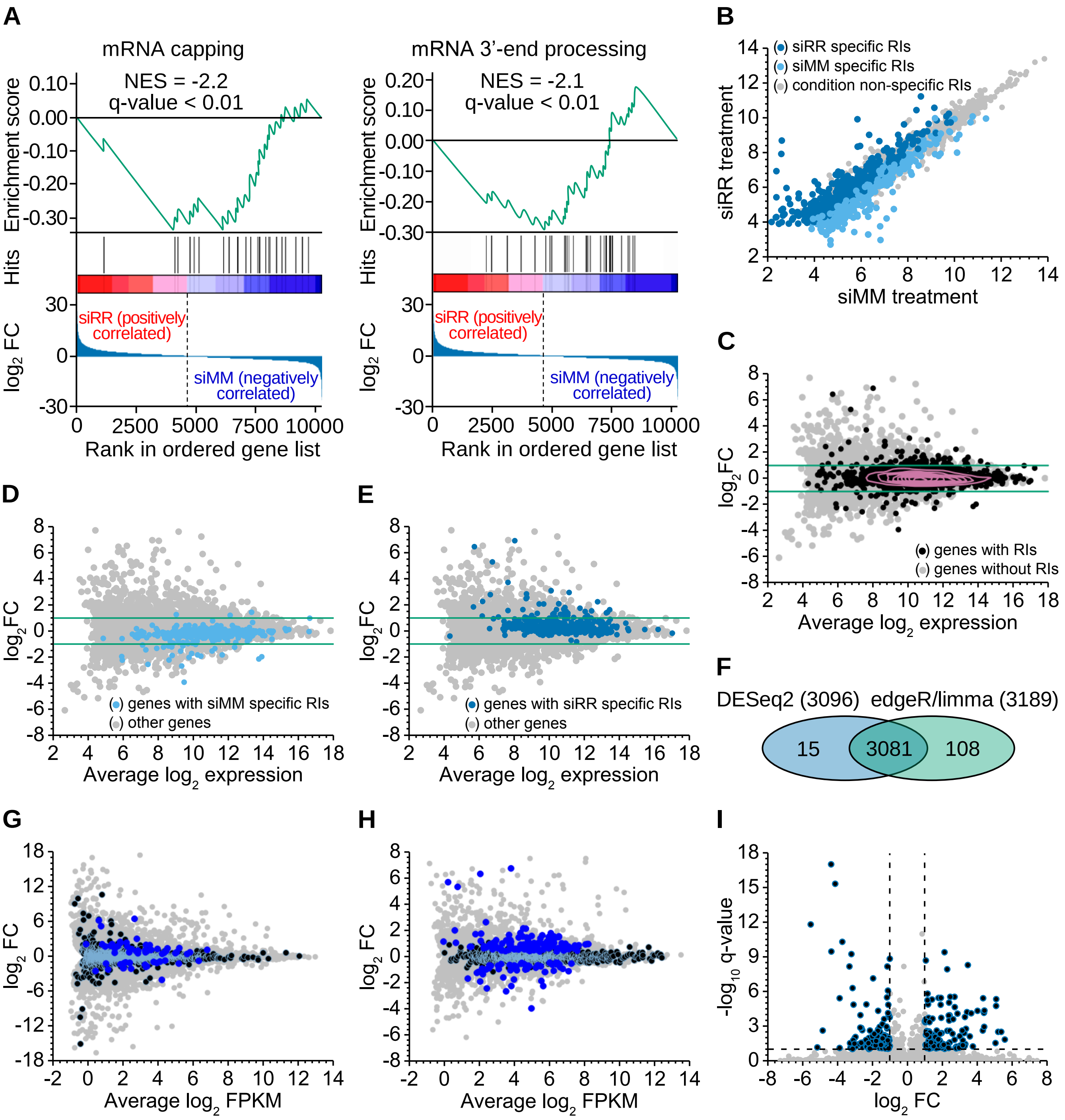


**Figure S1.** Knockdown of the fusion oncogene *RUNX1/RUNX1T1* leads to differential usage of a subset of exons in the transcriptome of Kasumi-1 cells.

(A) The GSEA enrichment plots for the Reactome mRNA capping and mRNA 3'-end processing pathways. The normalized enrichment score and its statistical significance are shown in the each enrichment plot.

(B) Expression of the condition-specific RIs in the siRR- and siMM-treated leukemia cells. The axes denote the normalized and $log_2$ transformed expression values averaged over the three biological repeats. Here and below, siRR-specific RIs are the RIs with statistically significant expression in the siRR-treated but not in the siMM-treated leukemia samples. Similarly, siMM-specific RIs are the RIs with statistically significant expression only in the siMM-treated leukemia samples.

(C) to (E) Series of diagnostic MA-plots of gene expression under the two siRNA treatment conditions. The genes that produce the RIs form relatively stable core of distribution. This core is bounded by a density contour in (C). However, the genes that produce condition-specific RIs show a small but noticeable difference in expression. In these plots, bluish green lines indicate two-fold change in gene expression.

Parts B to E are based on the analysis of the RIs detected using the DESeq2 algorithm. Very similar results were observed for the RIs detected by the edgeR/limma algorithm.

(F) Diversity of the transcripts containing RIs detected using different approaches. For each approach, the total number of transcripts with RIs is indicated in parentheses.

(G) Diagnostic MA-plot of three groups of transcripts: i) –RIs transcripts (●), ii) +RIs transcripts without significant change in expression (●), and iii) +RIs transcripts with significant change in expression (●).

(H) Diagnostic MA-plot of three groups of genes: i) genes that produce only –RIs transcripts (●), ii) stably expressed genes which produce +RIs transcripts (●), and iii) differentially expressed genes which produce +RIs transcripts (●).

(I) The JunctionSeq algorithm identified 221 diffUEs distributed over 142 individual genes.

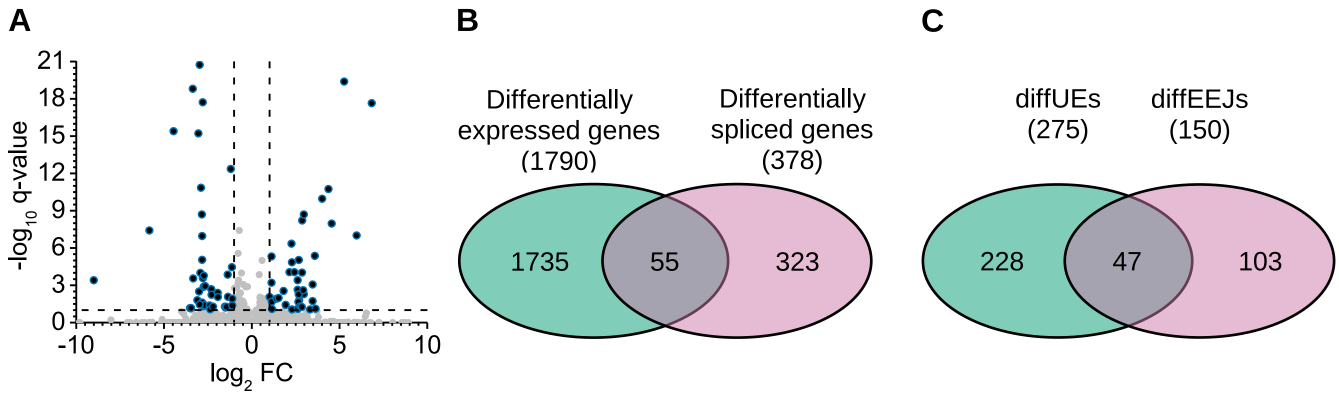


**Figure S2.** The knockdown of the fusion oncogene *RUNX1*/*RUNX1T1* leads to differential expression and differential splicing in a significant fraction of the genes in Kasumi-1 cells.

(A) The JunctionSeq algorithm identified 83 diffEEJs distributed over 70 individual genes.

(B) Of all the genes with differential splicing, only 55 genes demonstrate differential expression.

(C) Of all the expressed genes, 378 genes demonstrate differential splicing following the *RUNX1/RUNX1T1* knockdown. Herewith, the diffUEs only was detected in 228 genes and the diffEEJs without diffUEs were identified in 103 genes. In addition, both types of differential splicing occur in 47 genes.

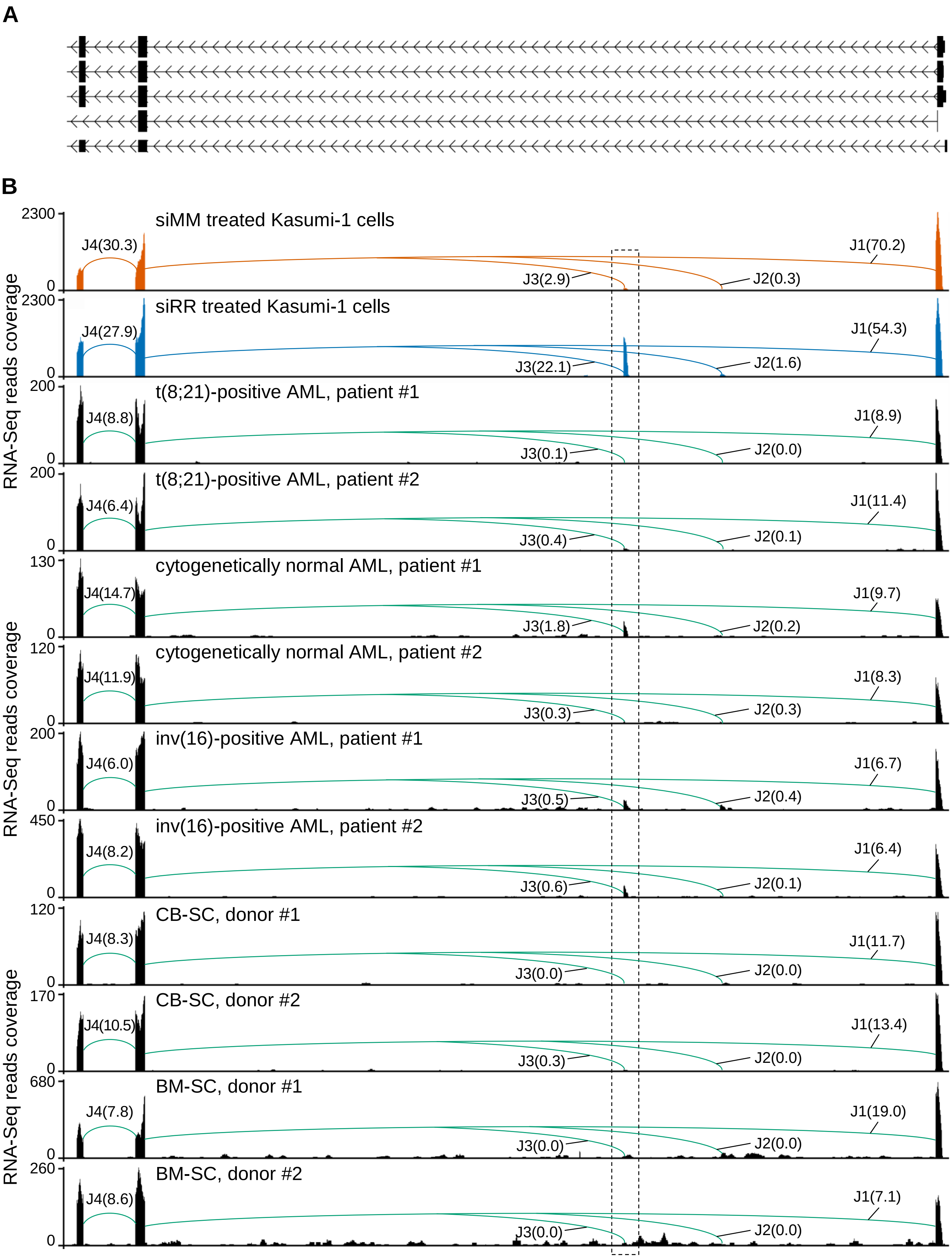


**Figure S3.** Splicing events at the 5'-end of *PARL* gene.
(A) Genomic location and organization of the 5' part of *PARL* gene.

(B) Splicing graphs of the 5’ part of *PARL* gene in the t(8;21)-positive AML cells, non-t(8;21)-positive AML cells, normal CB-SC and normal BM-SC. These graphs were reconstructed from the RNA-Seq reads mapped to the exons and EEJs of *PARL* gene. EEJs are designated by the letter J and numbered. The normalized number of reads supporting the identified junctions is shown in parentheses. The Region of interest is boxed.

A

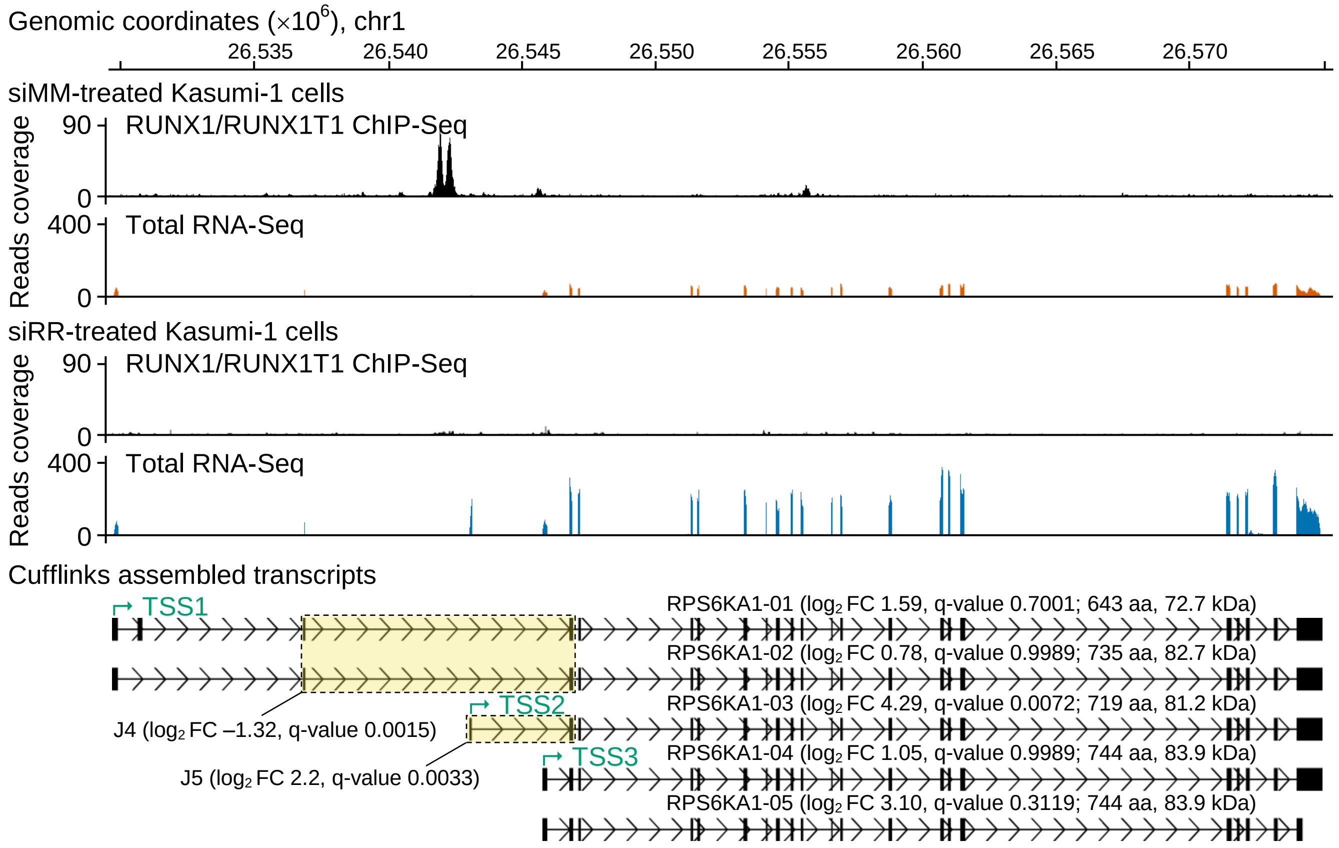

Genomic coordinates (×10^6), chr1
26,535
26,540
26,545
26,550
26,555
26,560
26,565
26,570
siMM-treated Kasumi-1 cells
Reads coverage
90
0
RUNX1/RUNX1T1 ChIP-Seq
400
0
Total RNA-Seq
siRR-treated Kasumi-1 cells
Reads coverage
90
0
RUNX1/RUNX1T1 ChIP-Seq
400
0
Total RNA-Seq
Cufflinks assembled transcripts
TSS1
RPS6KA1-01 (log2 FC 1.59, q-value 0.7001; 643 aa, 72.7 kDa)
RPS6KA1-02 (log2 FC 0.78, q-value 0.9989; 735 aa, 82.7 kDa)
TSS2
RPS6KA1-03 (log2 FC 4.29, q-value 0.0072; 719 aa, 81.2 kDa)
J4 (log2 FC –1.32, q-value 0.0015)
TSS3
RPS6KA1-04 (log2 FC 1.05, q-value 0.9989; 744 aa, 83.9 kDa)
J5 (log2 FC 2.2, q-value 0.0033)
RPS6KA1-05 (log2 FC 3.10, q-value 0.3119; 744 aa, 83.9 kDa)


B

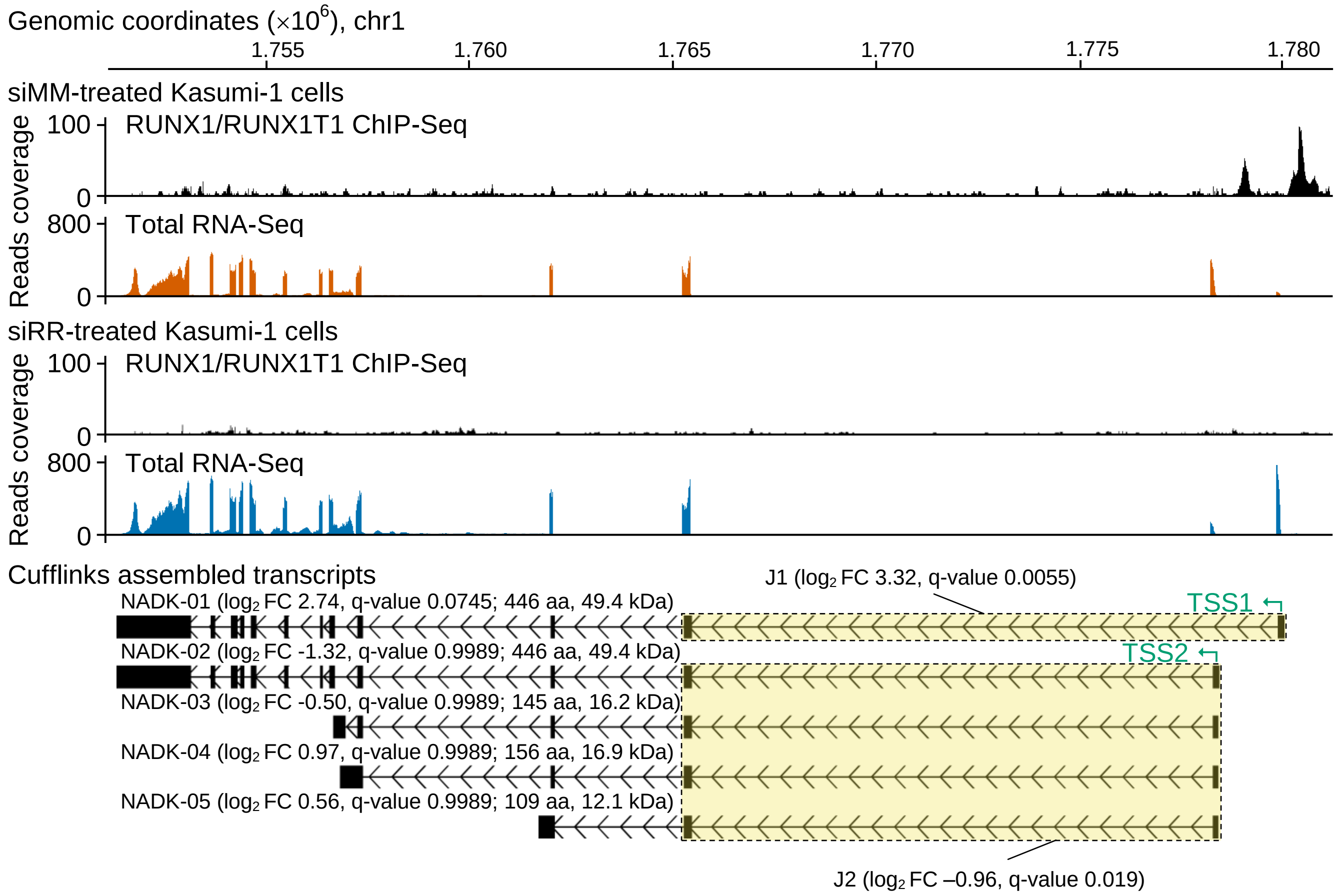

Genomic coordinates (×10^6), chr1
1.755
1.760
1.765
1.770
1.775
1.780
siMM-treated Kasumi-1 cells
Reads coverage
100
0
RUNX1/RUNX1T1 ChIP-Seq
800
0
Total RNA-Seq
siRR-treated Kasumi-1 cells
Reads coverage
100
0
RUNX1/RUNX1T1 ChIP-Seq
800
0
Total RNA-Seq
Cufflinks assembled transcripts
J1 (log2 FC 3.32, q-value 0.0055)
NADK-01 (log2 FC 2.74, q-value 0.0745; 446 aa, 49.4 kDa)
TSS1
NADK-02 (log2 FC -1.32, q-value 0.9989; 446 aa, 49.4 kDa)
TSS2
NADK-03 (log2 FC -0.50, q-value 0.9989; 145 aa, 16.2 kDa)
NADK-04 (log2 FC 0.97, q-value 0.9989; 156 aa, 16.9 kDa)
NADK-05 (log2 FC 0.56, q-value 0.9989; 109 aa, 12.1 kDa)
J2 (log2 FC –0.96, q-value 0.019)

**Figure S4.** Knockdown of the fusion oncogene *RUNX1/RUNX1T1* leads to differential splicing of *RPS6KA1* (A) and *NADK* (B) transcripts in Kasumi-1 cells.

In this figure, each gene specific panel includes genomic coordinates, read coverage tracks and a set of Cufflinks-assembled full-length transcripts. Read coverage tracks demonstrate the RUNX1/RUNX1T1 ChIP-Seq and total RNA-Seq results separately for siMM- and siRR-treated Kasumi-1 cells. Cufflinks transcripts are provided with Cuffdiff-based $\log_2$ FC and q-values as well as the size of *in silico* predicted proteins. The positions of the Cuffdiff-determined transcription start sites are also shown using green arrows. In addition, diffEEJs are highlighted in yellow and are accompanied by statistics obtained using the diffSplice algorithm.

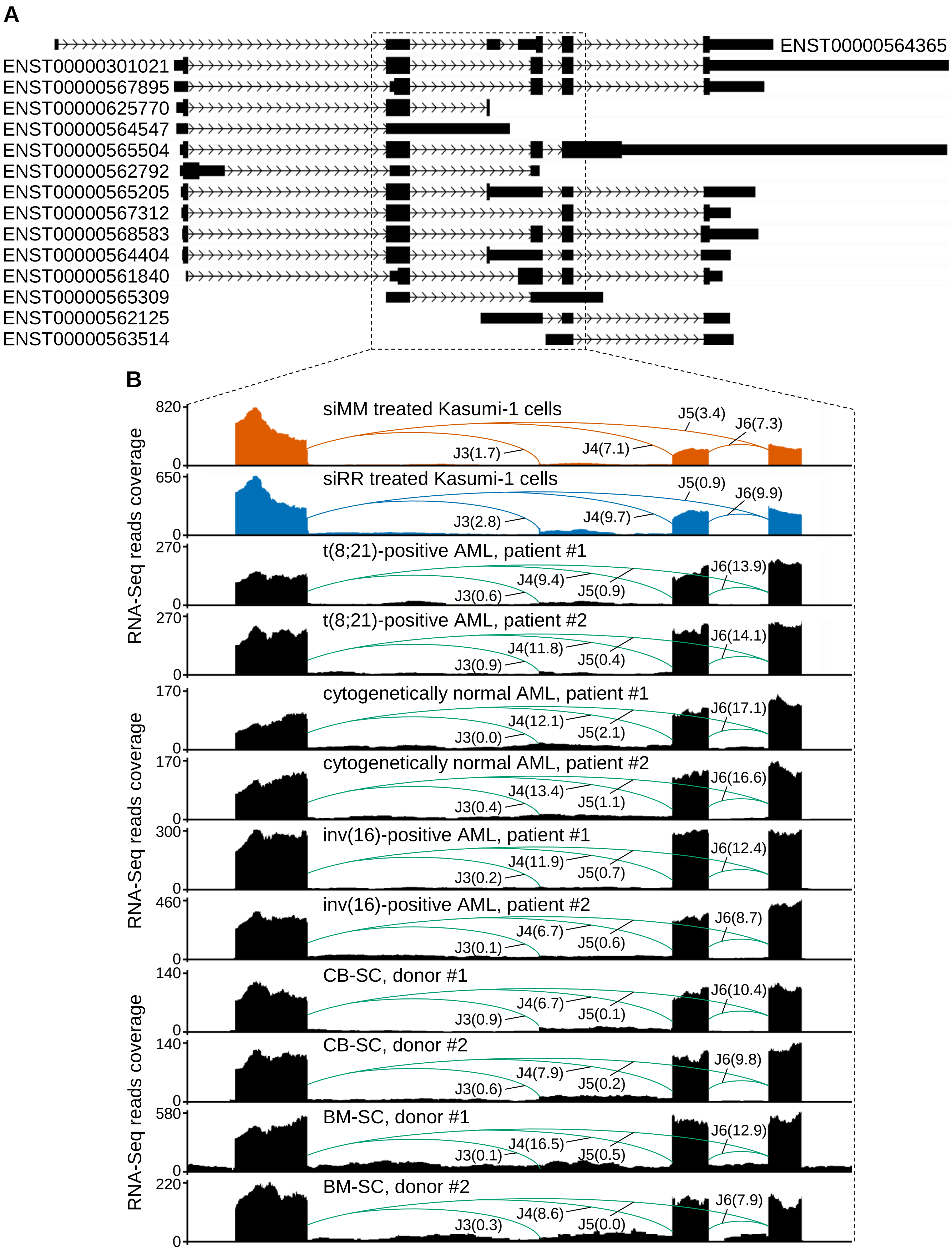


**Figure S5.** Splicing events at the middle part of *TRAPPC2L* gene.
(A) Genomic location and organization of *TRAPPC2L* gene. The middle part of the gene is

boxed. In this part of the gene, there can be alternative termination of transcription, alternative transcription start site, a selection of alternative 5' and/or 3' splice sites, and cassette exons.

(B) Splicing graphs of the middle part of *TRAPPC2L* gene in the t(8;21)-positive AML cells, non-t(8;21)-positive AML cells and normal CD34-positive hematopoietic stem/progenitor cells. These graphs were reconstructed from the RNA-Seq reads mapped to the exons and EEJs of *TRAPPC2L* gene. EEJs are designated by the letter J and numbered. The normalized number of reads supporting the identified junctions is shown in parentheses.

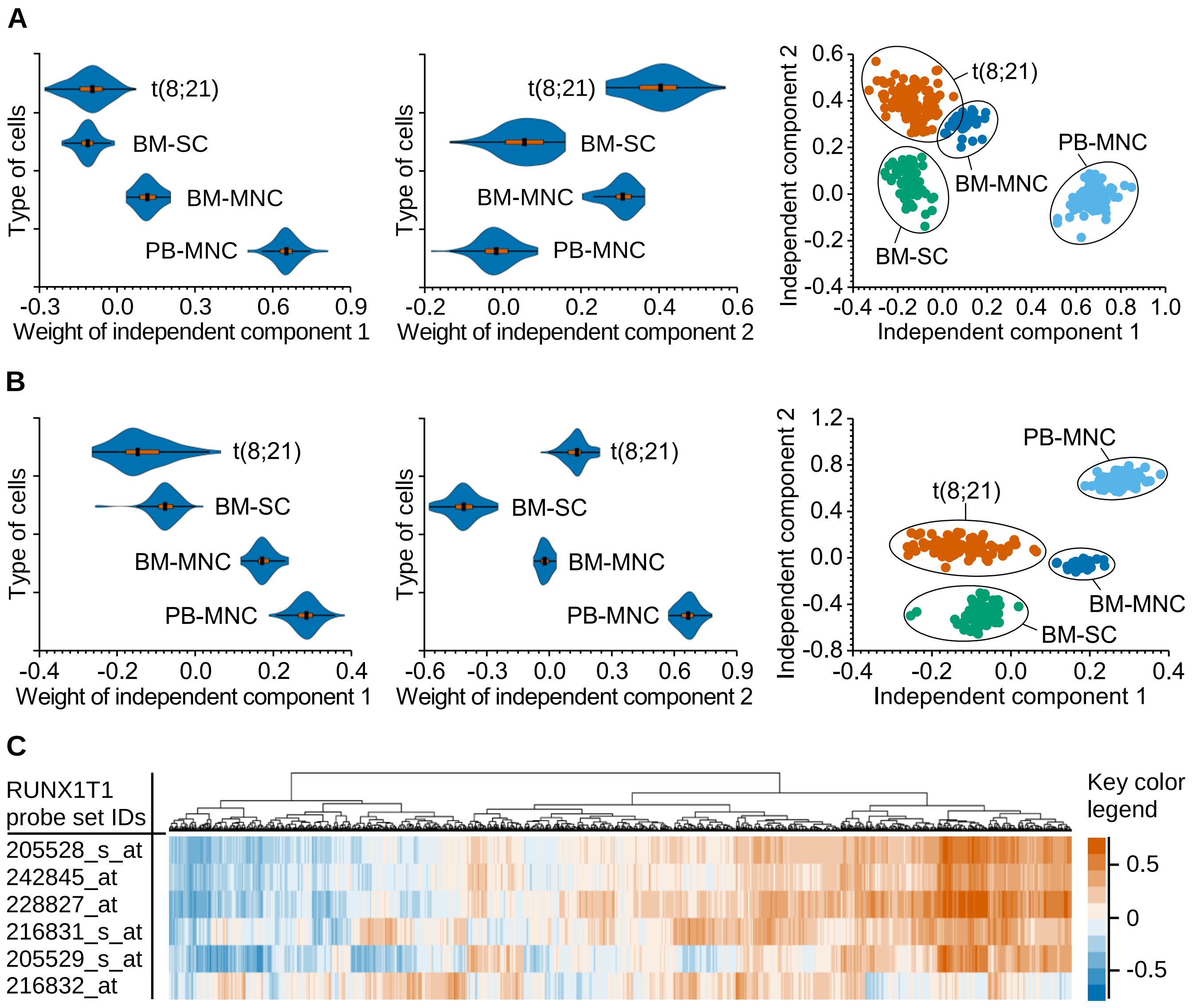


**Figure S6.** Independent component analysis identified a leukemia specific signature in the expression of genes encoding splicing factors and mRNA surveillance genes. This analysis was based on the public microarray data collected from 106 t(8;21)-positive AML samples (designated as t(8;21)), 85 samples of normal BM-SC, 32 samples of normal bone marrow mononuclear cells (BM-MNC) and 100 samples of normal peripheral blood mononuclear cells (PB-MNC) using Affymetrix® HG-U133 Plus 2.0 gene chip. Moreover, the publicly available RNA-Seq data for 20 t(8;21)-positive AML primary samples were also included in the correlation analysis.

(A) Independent component analysis of the whole set of genes expressed in the samples of interest. This type of analysis revealed two subsets of genes that distinguish the t(8;21)-positive leukemia cells from the normal cells of hematopoietic origin (right scatter plot). These genes form independent component 1 (1681 genes, FDR $< 0.05$; left violin plot) and independent component 2 (1071 genes, FDR $< 0.05$; middle violin plot). One-way ANOVA test confirms the inequality of the means between the different types of cells (p-value $< 0.001$ for each of independent components). Moreover, according to the two-sided Fisher's exact test, each of the components is statistically significantly (p-value $< 0.001$) enriched (1.62 and 1.66 fold enrichment, respectively, for independent component 1 and independent component 2) with genes coding splicing factors and mRNA surveillance genes. In total, these independent components contain 150 genes encoding splicing factors and mRNA surveillance genes.

(B) Independent component analysis of only a subset of splicing factor genes and mRNA surveillance genes expressed in the samples of interest. According to this type of analysis, t(8;21)-positive leukemia cells can be clearly separated from normal cells of hematopoietic origin using only expression data of the genes encoding splicing factors and mRNA surveillance genes (right scatter plot). It is noteworthy, that not all the genes encoding splicing factors and mRNA surveillance genes are required for this, but only two subsets of such genes that form two new independent components (violin plots).

(C) Heatmap of the co-expression of *RUNX1*/*RUNX1T1* and the genes encoding splicing factors and mRNA surveillance genes in 106 t(8;21)-positive AML samples. The co-expression was inferred from microarray data by calculating the Pearson's correlation coefficient. In this co-expression analysis, the RUNX1T1 probe sets (left side of picture, rows) were used as specific indicators of *RUNX1*/*RUNX1T1* expression because *RUNX1T1* gene itself is not active in the t(8;21)-positive leukemia cells. The upper dendrogram (columns) represents the results of hierarchical clustering of genes encoding splicing factors and mRNA surveillance genes. Key color legend shows the range of values of the Pearson's correlation coefficient.
